# Dual-targeted nanoparticles enable the ex vivo generation of cytotoxic and phagocytic CAR effector cells

**DOI:** 10.64898/2026.09.21.753074

**Authors:** Jisu Hong, Jinwoong Lim, Minho Park, Jimin Song, Byung Joon Lee, Miyoung Park, Min-Kyoo Shin, Hyungseok Seo, Hyoung Jin Kang, Byung-Soo Kim, Chang-Han Lee

## Abstract

Chimeric antigen receptor (CAR)-T cells and CAR-macrophages (or CAR-monocytes) offer complementary antitumor functions. CAR-T cells exert potent cytotoxicity against cancer cells, whereas CAR-macrophages can infiltrate solid tumors, phagocytose cancer cells and promote antigen spreading. Antibody-conjugated lipid nanoparticles (LNPs) have emerged as non-viral mRNA delivery platforms for generating CAR-immune cells. However, these systems have generally been optimized to target a single immune-cell type. Here, to enable the ex vivo generation of CAR-T cells and CAR-monocytes, we developed CLCM08, an anti-CD222 antibody, and co-displayed it with an anti-CD5 antibody on LNPs. CD222 is predominantly expressed by monocytes, macrophages and T cells, and facilitates the internalization of CD222-binding molecules. CD5 is primarily expressed by T cells and has previously been used to enable LNP-mediated mRNA delivery to these cells. Delivery of CAR mRNA by dual CD222/CD5 antibody-conjugated LNPs to primary monocytes and T cells substantially enhanced the ex vivo generation of CAR-monocytes and CAR-T cells compared with non-conjugated LNPs. Moreover, CD19 CAR-T cells and CAR-monocytes generated using dual CD222/CD5 antibody-conjugated LNPs showed enhanced cytotoxic and phagocytic activity, respectively, relative to cells generated using non-conjugated LNPs, with anti-CD222 LNPs producing the largest phagocytic gain in monocytes. These findings establish an adaptable antibody-conjugated LNP-mRNA platform for the ex vivo generation of CAR-T cells and CAR-monocytes.

**Highlights:** • Dual CD222/CD5-targeted LNPs enhanced CAR mRNA delivery to monocytes and T cells.

• Dual-targeted LNPs improved ex vivo generation of CAR-monocytes and CAR-T cells.

• CD19 CAR-T cells generated with dual-targeted LNPs showed enhanced cytotoxicity.

• Dual- and CD222-targeted LNPs both enhanced CAR-monocyte phagocytosis.

• Dual-targeted LNPs generated CAR effectors with complementary antitumor functions.

## 1. Introduction

Chimeric antigen receptor (CAR)-T cell therapy has revolutionized the treatment of hematological malignancies [1]. However, its efficacy against solid tumors remains limited by the immunosuppressive tumor microenvironment (TME), tumor heterogeneity, and inefficient tumor infiltration [2,3]. CAR-engineered myeloid cells, including macrophages and their circulating precursors, monocytes, offer complementary properties that may help overcome these barriers [4,5]. Whereas CAR-T cells provide rapid and potent cytotoxicity against target cancer cells [6], CAR-monocytes leverage their intrinsic trafficking properties to infiltrate dense solid tumors and differentiate into CAR-macrophages within the TME [3,7,8]. Once differentiated, these cells can phagocytose cancer cells [4,5] and promote broader antitumor immunity through antigen processing, cross-presentation [4,9], and antigen spreading [10]. Thus, the combination of lymphoid and myeloid CAR-immune cells may provide a complementary therapeutic strategy that integrates potent cytotoxicity with efficient tumor infiltration, phagocytosis, and immune amplification.

Despite this potential, the concurrent generation of distinct CAR-engineered immune-cell populations remains technically challenging. Conventional ex vivo CAR-cell manufacturing relies predominantly on viral vectors, which entail high production costs, complex manufacturing workflows and concerns regarding genomic integration and manufacturing scalability [11–13]. Moreover, although viral transduction is well established in activated T cells, it is considerably less efficient and less readily applicable to monocytes [4,14,15]. Lipid nanoparticles (LNPs) have therefore emerged as attractive non-viral platforms for the transient delivery of CAR mRNA, offering a potentially safer and more flexible approach to CAR-cell engineering [8,13,16,17]. Surface functionalization of LNPs with antibodies or other targeting ligands [18,19] can further enable selective mRNA delivery to defined immune cell populations [20,21]. However, current targeted LNP platforms have been largely developed to recognize and transfect a single immune-cell lineage [8,20–22]. This single-lineage bias presents a fundamental limitation for the simultaneous engineering of lymphoid and myeloid cells from ex vivo multiple-lineage cells, necessitating separate targeting and manufacturing processes for combination cellular therapies [8,20].

To overcome this limitation, we sought to develop a dual-targeting LNP platform capable of simultaneously engaging both T cells and monocytes. CD5 is a well-characterized surface glycoprotein predominantly expressed by T cells [23,24], and antibody-mediated targeting of CD5 has previously been shown to facilitate LNP-mediated CAR mRNA delivery to T cells [20,25]. As a complementary target for myeloid-cell engagement, we focused on CD222, also known as the cation-independent mannose-6- phosphate receptor (CI-MPR) [26]. CD222 is abundantly expressed on monocytes and macrophages [27] and is also present on T cells [28]. Importantly, CD222 undergoes rapid receptor-mediated internalization upon ligand binding, providing a favorable route for LNP uptake and intracellular delivery [26,29–31]. We therefore hypothesized that simultaneous engagement of CD5 and CD222 could broaden the cellular tropism of LNPs and enable coordinated CAR mRNA delivery across lymphoid and myeloid lineages.

Here, we develop an adaptable dual-targeted LNP platform for the simultaneous ex vivo generation of CAR-T cells and CAR-monocytes. We generated CLCM08, a novel anti-CD222 antibody, and co-displayed it with an anti-CD5 antibody on the surface of CAR mRNA-loaded LNPs. These dual-targeted LNPs efficiently delivered CAR mRNA to primary T cells and monocytes, substantially enhancing the concurrent generation of CAR-T cells and CAR-monocytes compared with non-targeted LNPs. Using a CD19-targeting CAR as a model system, we further demonstrate that the resulting CAR-immune cells retain complementary effector functions, combining potent T cell-mediated cytotoxicity with efficient monocyte-mediated phagocytosis. Collectively, these findings establish a single-pot, LNP–mRNA-based platform for the coordinated engineering of complementary CAR-immune cell populations, providing a versatile strategy for the development and clinical translation of multi-cellular immunotherapies against solid tumors.

## 2. Materials and methods

### 2.1 Cell lines and culture conditions

THP-1, Jurkat, Raji, and SNU-5 cells were obtained from the American Type Culture Collection (ATCC). THP-1, Jurkat, Raji and SNU-5 cells were maintained in RPMI-1640 supplemented with 10% heat-inactivated fetal bovine serum and 1% antibiotic-antimycotic solution. RAW 264.7 cells were maintained in DMEM high glucose media supplemented with 10% heat-inactivated fetal bovine serum and 1% antibiotic-antimycotic solution. THP-1 cultures were supplemented with 0.05 mM 2-mercaptoethanol when required. For yeast-display cell-based enrichment, THP-1 cells were differentiated with 100 nM phorbol 12-myristate 13-acetate (PMA) for 24 h and subsequently rested in PMA-free medium for 24 h before fixation [32]. Cells were maintained at 37 °C in 5% CO2 and confirmed to be mycoplasma-free using the e-Myco™ Mycoplasma PCR Detection Kit (ver. 2.0; Cat. No. 25235; iNtRON Biotechnology, Seongnam, Gyeonggi-do, Republic of Korea).

### 2.2 Human PBMCs and ethical approval

Clinical leukapheresis products were obtained from healthy adult volunteers under a protocol approved by the Institutional Review Board of Seoul National University Hospital (SNUH-IRB 1606-033-768), with written informed consent. Peripheral blood mononuclear cells (PBMCs) were isolated from apheresis products without delay and processed within 24 h of receipt. Isolated PBMCs were cryopreserved in aliquots and stored in liquid nitrogen until use, so that all experiments could be performed from a common cell bank. For each experiment, frozen PBMCs were thawed rapidly, diluted into pre-warmed complete RPMI-1640, centrifuged at 300 × g for 3 min, and recovered overnight in complete medium containing 50 U/mL human IL-2 (Enzynomics, Daejeon, Republic of Korea). Unless otherwise indicated, PBMCs recovered under these conditions without subsequent CD3/CD28 stimulation are referred to throughout as non-activated PBMC cultures. All primary PBMC and T-cell experiments were performed using cells derived from the same donor, and replicate measurements represent technical replicates of that donor.

### 2.3 Public single-cell RNA-sequencing analysis

Public single-cell RNA-sequencing datasets were analyzed to assess expression of IGF2R, the gene encoding CD222, across circulating immune-cell populations. A pan-cancer peripheral blood immune-cell single-cell atlas was curated from nine publicly available scRNA-seq datasets (GSE139324 [33], CancerSCEM_GC-035 [34], GSE155698 [35], GSE114725 [36], GSE145281 [37], CancerSCEM_GBM-108 [34], CancerSCEM_BRCA-071 [34], GSE242780 [38], and GSE117988 [39]), comprising 248,266 peripheral blood-derived cells across 77 sample-level profiles. Dataset accessions and preprocessing details are provided in **Supplementary Table S1**.

The curated datasets were concatenated into a unified AnnData object and processed using Scanpy [40]. Raw counts were normalized to a target sum of 10,000 counts per cell using sc.pp.normalize_total() and log-transformed using sc.pp.log1p(). Highly variable genes were identified using sc.pp.highly_variable_genes() with dataset accession specified as the batch key, followed by scaling and principal-component analysis. Harmony integration [41] was performed in principal-component space using dataset accession as the batch covariate. A neighborhood graph was constructed from the Harmony-corrected representation using 30 nearest neighbors, followed by UMAP dimensionality reduction. Initial cell-type annotations were assigned using SCimilarity [42] predictions and subsequently reviewed and manually corrected based on canonical lineage-marker expression and the local neighborhood structure of the integrated embedding.

IGF2R expression was visualized using UMAP feature plots, violin plots, and dot plots. Within each cell type, IGF2R-high cells were defined as the top 20% of cells ranked by IGF2R expression, whereas the remaining cells were classified as IGF2R-non-high. Raw counts were aggregated into donor-level pseudobulk profiles for each donor, cell type, and IGF2R state. Differential-expression analysis was performed separately for each cell type using PyDESeq2 [43] with a paired donor-level model. IGF2R- high samples were compared with donor-matched IGF2R-non-high samples using the DESeq2 Wald test, with P values adjusted using the Benjamini–Hochberg procedure. Genes with an adjusted P value < 0.05 and an absolute log_2_ fold change ≥ 0.5 were considered differentially expressed.

For pathway analysis, all tested genes were ranked according to the DESeq2 Wald test statistic, and preranked gene set enrichment analysis was performed using GSEApy [44] against gene-set collections from the Kyoto Encyclopedia of Genes and Genomes (KEGG 2026) and the Molecular Signatures Database (MSigDB Hallmark 2020). Gene sets containing fewer than 15 or more than 500 genes were excluded, and pathways with an FDR q value ≤ 0.05 were considered significantly enriched and prioritized for biological interpretation; non-significant enrichments, where shown, were presented descriptively.

### 2.4 Recombinant human CD222 preparation

A DNA sequence encoding residues 36–466 of human CD222/CI-MPR was cloned into pcDNA3.4 with a C-terminal hexahistidine tag. Recombinant protein was expressed in Expi293F cells, purified from clarified supernatant by Ni-NTA chromatography, and buffer-exchanged into PBS. Purity was assessed by SDS-PAGE and concentration by absorbance at 280 nm. Purified CD222-His was biotinylated using Sulfo-NHS-LC-Biotin (Thermo Fisher Scientific, Waltham, MA, USA) according to the manufacturer’s instructions, including removal of excess unreacted biotin following the labeling reaction.

### 2.5 Mouse immunization and serum titer

Six-week-old female C57BL/6 mice (n = 4) were immunized intraperitoneally with 50 μg of recombinant CD222-His formulated at a 1:1 volume ratio with Imject™ Alum Adjuvant (Thermo Fisher Scientific). Booster immunizations were administered every 2 weeks at weeks 2, 4, 6, and 8. Blood was collected weekly for serum-titer measurement. All animal procedures were approved by the Seoul National University Institutional Animal Care and Use Committee (SNU-220406-6-2). For serum ELISA, plates were coated with 100 ng of CD222-His per well, blocked with 3% BSA, incubated with serially diluted serum, and developed using HRP-conjugated goat anti-mouse IgG and TMB substrate.

### 2.6 Immune Fab yeast-display library construction and screening

After the final immunization, total RNA was isolated from spleen using QIAzol Lysis Reagent (QIAGEN, Hilden, Germany), and cDNA was generated using the SuperScript™ III First-Strand Synthesis System (Thermo Fisher Scientific). VH and VL genes were amplified, fused to human CH1 and CL regions by overlap-extension PCR, and introduced into JAR200 and YVH10 yeast strains. The haploid libraries were paired by mating, and Fab display was induced in 2× SGCAA medium for 48 h at 20 °C [45,46]. The library was enriched over three sequential rounds: magnetic-activated cell sorting (MACS) in round 1 and fluorescence-activated cell sorting (FACS) in round 2, both using 1 μM biotinylated CD222-His for labeling, followed by cell-based biopanning against fixed PMA-differentiated THP-1 cells in round 3. Cell-based enrichment was performed using fixed PMA-differentiated THP-1 cells. Individual clones from the final enriched pool were sequenced, yielding 20 unique CD222-reactive clones designated CLCM01–CLCM20.

### 2.7 Recombinant antibody expression and purification

Selected mouse variable regions were reformatted as human IgG1/κ chimeric antibodies and transiently expressed in Expi293F cells. Supernatants were harvested after 5 days, and antibodies were purified using PuriOse™ Protein A FF resin (Amicogen, Jinju, Gyeongsangnam-do, Republic of Korea). Antibodies were buffer-exchanged into PBS, quantified by absorbance at 280 nm, and evaluated by SDS-PAGE and SEC-HPLC. Five clones were carried forward for full characterization, and CLCM08 was selected as the lead clone.

### 2.8 Anti-CD5 and control antibodies

The anti-human CD5 antibody used for LNP conjugation was the H65 clone, which was recombinantly expressed and purified in-house as a human IgG1/κ antibody. The S309 clone, also recombinantly expressed and purified in-house as a human IgG1/κ antibody, was used as an irrelevant isotype control where indicated. Antibodies were buffer-exchanged into PBS before DBCO modification.

### 2.9 Recombinant-antigen and cell-surface binding

For direct ELISA, immobilized CD222-His was incubated with serially diluted anti-CD222 IgG and detected with HRP-conjugated anti-human IgG. For cell-surface binding, THP-1 cells, Jurkat cells, or PBMCs were incubated with biotinylated anti-CD222 antibodies at 4 °C after Fc-receptor blocking with an in-house-produced human IgG1 isotype control antibody. Bound antibody was detected using Streptavidin, R-Phycoerythrin Conjugate (SAPE; Cat. No. S866; Thermo Fisher Scientific). PBMC monocytes were identified as CD14-positive cells and T cells as CD3-positive cells, with viability and singlet gating performed as shown in **Supplementary Fig. 5**.

### 2.10 Activation-dependent CD222 surface expression

PBMCs or isolated CD3-positive T cells were analyzed before and after CD3/CD28 activation at 24 h. For PBMC analysis, monocytes were identified using APC-conjugated anti-human CD14 antibody (Cat. No. 367118; BioLegend, San Diego, CA, USA), whereas T cells were identified using APC-conjugated anti-human CD3 antibody (Cat. No. 317318; BioLegend). CD222 surface binding in each population was assessed using the biotinylated anti-CD222 antibodies prepared as described above, followed by detection with Streptavidin, R-Phycoerythrin Conjugate (SAPE; Cat. No. S866; Thermo Fisher Scientific). Cells were analyzed by flow cytometry.

### 2.11 Temperature-dependent loss of surface-associated antibody signal

PBMCs were pre-incubated with an Fc-receptor blocking antibody, labeled with biotinylated anti-CD222 antibody at 4 °C and detected with streptavidin-PE. Samples were divided between 4 °C and 37 °C conditions and analyzed at 0, 0.5, 1, and 2 h. Surface-signal loss was calculated as [1 − (MFI_37 °C_/MFI_4 °C_)] × 100 within CD14-positive monocytes and CD3-positive T cells. Because the assay did not use acid stripping, fluorescence quenching, or imaging to distinguish surface from intracellular antibody, it was interpreted as temperature-dependent loss of surface-associated signal.

### 2.12 SEC-HPLC

Purified antibodies (40 μg in 200 μL) were analyzed using a Superdex 200 Increase 10/300 GL column (Cytiva, Marlborough, MA, USA) on a Nexera lite system (Shimadzu Corporation, Kyoto, Japan). Separation was performed in PBS at a flow rate of 0.5 mL/min with UV detection at 280 nm. Monomeric, high-molecular-weight, and low-molecular-weight species were quantified by peak-area integration.

### 2.13 mRNA preparation and CAR construct

EGFP mRNA and anti-CD19 CAR mRNA were generated by in vitro transcription from linearized plasmid templates using T7 RNA polymerase with full substitution of uridine by N1-methylpseudouridine and co-transcriptional CleanCap AG capping. The 5′ and 3′ untranslated regions and poly(A) tail were encoded in the DNA template. The CAR construct comprised a CD8α signal peptide, an anti-CD19 scFv (FMC63), an extracellular MYC tag, a CD28 hinge, a CD28 transmembrane domain, a CD28 costimulatory domain, and a CD3ζ signaling domain. Double-stranded RNA impurities were removed by cellulose-based purification and residual dsRNA and mRNA integrity were assessed by agarose gel electrophoresis.

### 2.14 LNP formulation

SM-102, DSPC, cholesterol, DMG-PEG2000, and DSPE-PEG2000-azide were dissolved in ethanol at a molar ratio of 50:10:38:1.5:0.5 [47]. mRNA was diluted in 10 mM citrate buffer at pH 4.0. The aqueous and ethanol phases were combined at 3:1 ratio via hand mixing method at an mRNA-to-SM-102 mass ratio of 1:10. Formulations were buffer-exchanged and concentrated into phosphate buffered saline (PBS, pH 7.2). Batch volume was 400 μL, mRNA recovery was approximately 60%, formulations were stored at 4 °C, and used within 48 h of preparation.

### 2.15 Linker screening, antibody functionalization, and LNP conjugation

DBCO-NHS reagents bearing C2 (HY-42973, MedChemExpress), C4 (HY-115524, MedChemExpress), PEG2 (HY-151827, MedChemExpress), PEG4 (HY-140272, MedChemExpress), or PEG24 (HY-151835, MedChemExpress) spacers were compared under matched conjugation conditions in RAW 264.7 cells using a model IgG rather than the anti-CD222 or anti-CD5 targeting antibodies used elsewhere in this study. The C2 spacer produced the highest reporter signal and was used for all antibody-LNP conjugates reported in this study. Antibodies were modified with DBCO-NHS ester at an antibody-to-linker molar ratio of 1:10 for 2 h at room temperature in PBS (pH 7.2) at an antibody concentration of 2 mg/mL. Excess unreacted linker was removed using a Zeba dye and biotin-removal column. DBCO-modified antibodies were incubated with azide-bearing LNPs overnight at room temperature. The nominal antibody-input ratio is defined relative to the molar amount of DSPE-PEG2000-azide incorporated into the LNP formulation. Single-ligand and dual-ligand formulations were prepared using the same nominal total IgG input, with the dual formulation split between anti-CD222 and anti-CD5.

### 2.16 Antibody-input titration and physicochemical characterization

Anti-CD222 LNPs were prepared at nominal antibody-input ratios of 1%, 10%, and 100% and tested for EGFP mRNA delivery in Jurkat and THP-1 cells. Hydrodynamic diameter, polydispersity index (PDI), and zeta potential were measured using a Zetasizer Nano ZS (Malvern Panalytical). Particle morphology was assessed by Cryo-TEM (Talos L120C, FEI). mRNA concentration and encapsulation efficiency were measured by the Quant-iT RiboGreen assay with and without Triton X-100 disruption. Size, PDI, zeta potential and encapsulation efficiency for every formulation used in this study are tabulated in **Supplementary Table S3**.

### 2.17 Receptor-dependence controls

For productive-delivery competition, Jurkat cells were preincubated with excess soluble anti-CD222 IgG, anti-CD5 IgG, or both for 30 min before treatment with the corresponding targeted LNP at 0.3 μg/mL mRNA. Each soluble competitor was added at a 10-fold molar excess relative to the nominal amount of the corresponding antibody conjugated to the targeted LNP formulation. EGFP expression was measured 24 h after LNP treatment. The same competition strategy was applied to primary-cell preparations, including CD14-positive monocytes within PBMC cultures and isolated CD3-positive T cells that had been activated with CD3/CD28 Dynabeads for 24 h before LNP treatment. To separate antigen-directed targeting from Fcγ receptor-mediated particle uptake, an Fc-silenced LALA-PG variant of CLCM08 was expressed, conjugated to EGFP mRNA-LNPs under matched conditions, and compared with wild-type CLCM08-LNP and with an Fc-silenced isotype-control (S309) LNP in CD14-positive monocytes within PBMC cultures.

### 2.18 T-cell isolation and activation

CD3-positive T cells were isolated from PBMCs by negative magnetic selection using the MojoSort™ Human CD3 T Cell Isolation Kit (BioLegend). Isolated T cells were cultured in complete RPMI-1640 supplemented with 50 U/mL human IL-2 and activated for 24 h using Dynabeads™ Human T-Activator CD3/CD28 for T Cell Expansion and Activation (Gibco, Grand Island, NY, USA) at a bead-to-cell ratio of 1:1 before LNP treatment. A parallel aliquot of the isolated T cells was maintained in the same IL-2– containing medium without CD3/CD28 Dynabeads for 24 h and is referred to as isolated non-activated T cells.

### 2.19 LNP-mediated mRNA delivery and flow cytometry

Unstimulated PBMCs, isolated non-activated T cells, or isolated CD3/CD28-activated T cells were seeded in non-treated 96-well round-bottom plates at 9 × 10^4^ cells in 150 μL per well and treated with EGFP mRNA-LNP or CAR mRNA-LNP at final mRNA concentrations of 0.3, 0.5, or 1.0 μg/mL for 24 h. EGFP expression was measured directly by flow cytometry. Surface CAR expression was detected using an in-house-produced biotinylated anti-MYC antibody (clone 9E10), followed by Streptavidin, R-Phycoerythrin Conjugate (SAPE; Cat. No. S866; Thermo Fisher Scientific). Monocytes were identified as live singlet CD14-positive events, whereas T cells were identified as live singlet CD3-positive events. Positivity gates were established using appropriate unstained, mock-treated, fluorescence-minus-one, and/or isotype controls. Data were acquired using a BD FACSCanto II flow cytometer (BD Biosciences, San Jose, CA, USA) and analyzed using FlowJo version 7.6 (FlowJo LLC, Ashland, OR, USA).

### 2.20 Viability, phenotype, and cytokine assays

Cell viability was assessed 24 h after LNP treatment by Annexin V/propidium iodide staining. Monocyte immunophenotypes were evaluated by flow cytometry using Brilliant Violet 421™ anti-human CD80 antibody (clone 2D10, Cat. No. 305221; BioLegend), PE anti-human CD86 antibody (clone BU63, Cat. No. 374206; BioLegend), APC/Fire™ 750 anti-human HLA-DR antibody (clone LN3, Cat. No. 327023; BioLegend), and FITC anti-human CD206 antibody (clone 15-2, Cat. No. 321104; BioLegend). T-cell phenotypes were evaluated using Brilliant Violet 421™ anti-human CD25 antibody (clone BC96, Cat. No. 302629; BioLegend), PE anti-human CD69 antibody (clone FN50, Cat. No. 12-0699-42; Invitrogen, Thermo Fisher Scientific), and APC anti-human CD279 (PD-1) antibody (clone A17188A, Cat. No. 379207; BioLegend). In PBMC culture supernatants, IL-1β, IL-6, TNF-α, and IFN-γ were quantified using the corresponding ELISA MAX™ Deluxe Sets (BioLegend) according to the manufacturer’s instructions. In isolated activated T-cell cultures, TNF-α and IFN-γ were quantified.

### 2.21 Phagocytosis assay in PBMC-derived CD14-positive monocytes

PBMCs were treated with CAR mRNA-LNPs for 24 h and cocultured with pHrodo Red-labeled CD19- positive Raji cells for 2 h at 37 °C at nominal effector-to-target (E:T) ratios of 10:1, 5:1, and 1:1. E:T ratios were calculated based on the total number of PBMCs added as effector cells rather than the number of CD14-positive monocytes. Following coculture, live singlet CD14-positive monocytes were analyzed by flow cytometry. Phagocytosis was quantified as the percentage of pHrodo Red^+^ cells among total CD14-positive monocytes.

### 2.22 CAR-T cytotoxicity assay

Activated primary T cells were treated with CAR mRNA-LNPs for 24 h. Dynabeads were removed before coculture with calcein-AM-labeled CD19-positive Raji cells at effector-to-target (E:T) ratios of 1:1, 2:1, and 4:1 for 6 h. Percent specific lysis was calculated from calcein release as [(experimental release − spontaneous release)/(maximum release − spontaneous release)] × 100. Spontaneous release was determined from calcein-AM-labeled Raji cells cultured without effector cells, whereas maximum release was generated by complete target-cell lysis using an in-house-prepared lysis buffer containing 2% (v/v) Triton X-100, 1% (w/v) SDS, 100 mM NaCl, and 1 mM EDTA.

### 2.23 Target-induced cytokine secretion

CAR mRNA-LNP-treated PBMC cultures or CAR mRNA-LNP-treated activated T-cell preparations were cocultured with CD19-positive Raji cells at an effector-to-target (E:T) ratio of 1:1 for 24 h. A Raji-only control was included. Following coculture, culture supernatants were harvested. IL-1β, IL-6, TNF-α, and IFN-γ were quantified in PBMC–Raji cocultures, whereas TNF-α and IFN-γ were quantified in activated T-cell–Raji cocultures, using the corresponding ELISA MAX™ Deluxe Sets (BioLegend) according to the manufacturer’s instructions. Cytokine outputs from PBMC cultures were interpreted as mixed-cell responses.

### 2.24 Statistical analysis

Data are presented as mean ± SD. For immune-cell-line experiments, n refers to independent experiments, and comparisons were performed using one-way ANOVA followed by Tukey’s multiple-comparison test, with statistical significance defined as P < 0.05. All primary human PBMC and T-cell experiments were performed using cells derived from a single healthy donor, and replicate measurements represent technical replicates of that donor. Technical replicates were therefore not treated as independent biological observations, and no inferential statistical testing was applied to the primary-cell data; these replicates are used only to indicate measurement variability, and the corresponding comparisons are reported descriptively. Exact n values are indicated in the corresponding figure legends. For gene set enrichment analysis, permutation-based P values are reported as bounded values rather than zero.

## 3. Results

### 3.1 Discovery and selection of an anti-CD222 antibody for LNP targeting

Single-cell transcriptomic datasets spanning eight tumor types showed transcription of IGF2R, the gene encoding CD222, across immune-cell populations, with comparatively high expression in monocyte and granulocyte compartments (**Fig. 1a** and **Supplementary Fig. 1a**–**e**; **Supplementary Tables S1** and **S2**). An exploratory donor-level pseudobulk comparison of IGF2R-high and IGF2R-non-high cells is provided as contextual analysis in **Supplementary Fig. 1f**–**h**; because this analysis is descriptive and was not designed to resolve trafficking behavior, it is not used to support a mechanistic interpretation.

**Fig. 1.**
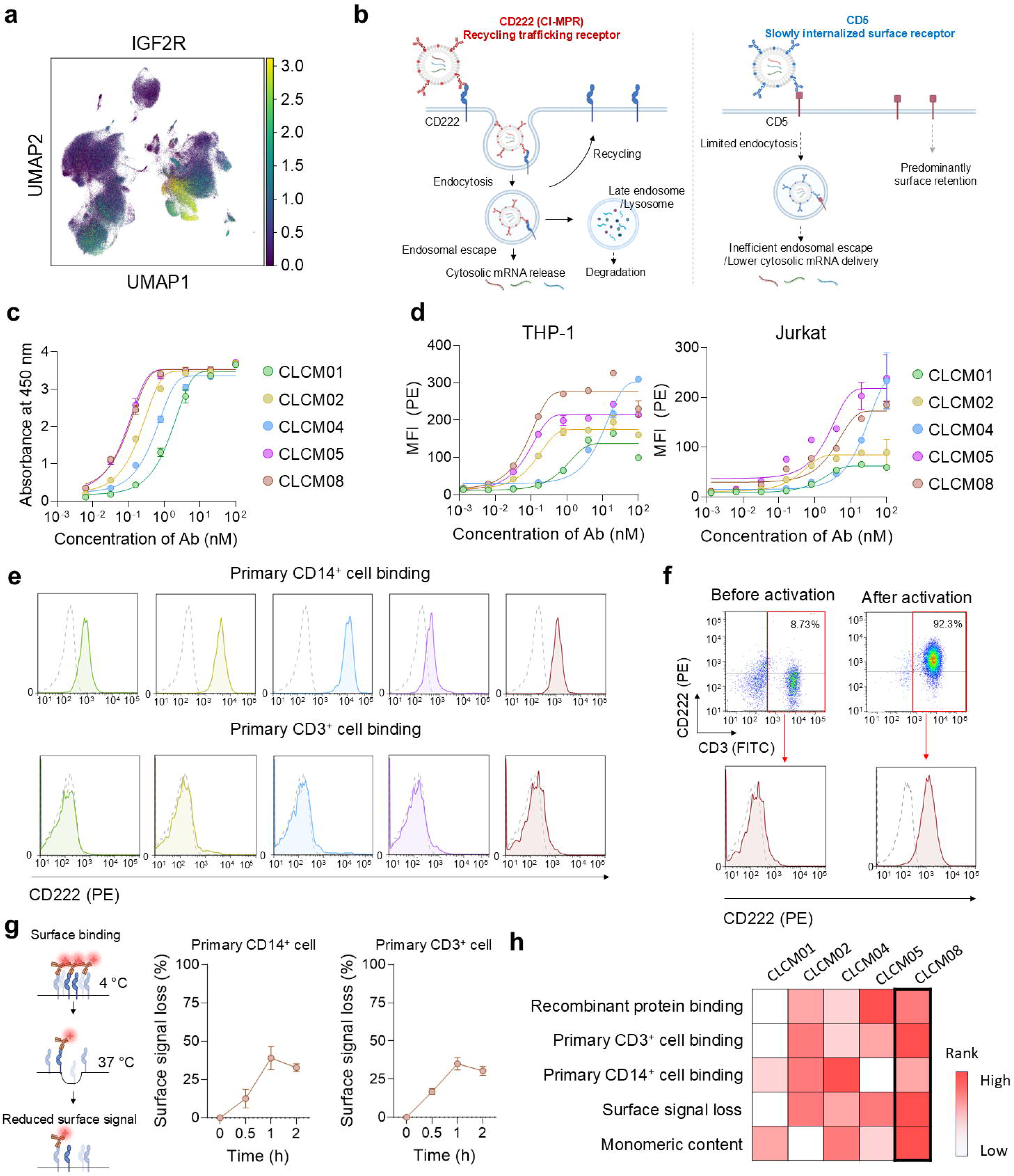
Discovery and selection of an anti-CD222 antibody for LNP targeting. (a) Uniform manifold approximation and projection of IGF2R (CD222) transcript abundance across immune-cell populations from public single-cell datasets. (b) Schematic of the two targeting axes compared in this study: CD222 undergoes constitutive surface-to-endosome trafficking, whereas antibody-bound CD5 is internalized more slowly. The schematic depicts the design rationale and is not intended to represent a measured intracellular routing or endosomal-escape mechanism. (c) Binding of the five candidate antibodies to immobilized recombinant CD222 by ELISA. (d) Binding to cell-surface CD222 on THP-1 and Jurkat cells. (e) Antibody staining of CD14-positive monocytes and CD3-positive T cells within non-activated PBMC cultures. (f) Representative CD222 surface staining of CD3-positive T cells before and after CD3/CD28 activation. (g) Loss of surface-associated antibody signal (%) after transfer from 4 to 37 °C in CD14- positive monocytes and CD3-positive T cells; the corresponding time courses for all five candidate clones are shown in Supplementary Fig. 6. (h) Summary comparison of the candidate antibodies across recombinant-antigen binding, primary-cell binding, temperature-dependent surface-signal loss, and monomeric content. Primary-cell data (panels e–g) are from a single donor; panel (g) is presented as mean ± SD of n = 2 technical replicates from that donor, and panels (e) and (f) show representative flow cytometry data. No inferential statistics were applied (Methods 2.24). Recombinant-antigen and cell-line data (panels c and d) are presented as mean ± SD from n = 2 independent experiments.

Based on this expression landscape and the reported trafficking properties of CD222, we defined the two targeting axes compared in this study [26,27]. CD222 was treated as an antibody-addressable trafficking receptor that cycles between the plasma membrane and endosomal compartments, whereas CD5 was treated as a lymphocyte-enriched surface receptor whose antibody-bound complexes are internalized slowly relative to CD222 [48]. This contrast in receptor availability, rather than a defined intracellular routing mechanism, provided the rationale for comparing CD222-, CD5-, and dual CD222/CD5-targeted LNP formulations across different immune-cell preparations (**Fig. 1b**).

To establish an antibody reagent capable of exploiting CD222 as a targeting receptor, we next generated and screened a panel of anti-CD222 antibodies. Mice were immunized with a recombinant N-terminal fragment of human CD222 spanning residues 36–466. Immune Fab yeast-display screening and sequential enrichment against recombinant CD222 and CD222-expressing cells yielded 20 CD222-reactive clones (**Supplementary Figs. 2–4**). Five clones, CLCM01, CLCM02, CLCM04, CLCM05, and CLCM08, were reformatted as human IgG1/κ chimeric antibodies and advanced for characterization.

We first examined whether the selected antibodies recognized both recombinant CD222 and the native receptor displayed on immune-cell surfaces. All five antibodies bound immobilized CD222 in a concentration-dependent manner and recognized cell-surface CD222 on THP-1 and Jurkat cells (**Fig. 1c,d**). On primary cells, all five antibodies showed increased staining of CD14-positive monocytes relative to the streptavidin-PE-only control, whereas staining of CD3-positive T cells exhibited low but detectable staining levels (**Fig. 1e** and **Supplementary Fig. 5**). After CD3/CD28 activation, the CD222- positive T-cell fraction increased from 8.73% to 92.3% in a representative sample (**Fig. 1f**). Because donor-level replication of this shift is not available in the current dataset, subsequent comparisons are described as preparation-associated rather than activation-specific. Accessible CD222 is therefore present on monocytes without stimulation and appears on T cells only after activation, which sets up the preparation-dependent comparison that follows.

To assess whether surface-bound anti-CD222 antibodies exhibited temperature-dependent behavior consistent with receptor uptake or trafficking, we monitored the loss of cell-associated fluorescence after warming. Warming antibody-labeled cells from 4 °C to 37 °C reduced surface-associated signal over time (**Fig. 1g** and **Supplementary Fig. 6**). Because the assay did not distinguish internalized antibody from shedding, masking, or other causes of signal loss, this readout is reported as temperature-dependent loss of surface-associated antibody rather than as internalization. On that measure, CLCM08 lost 39% of surface signal on monocytes and 35% on T cells at 1 h, indicating that the antibody-receptor complex does not remain statically displayed at the cell surface.

Finally, to select a lead antibody suitable for LNP conjugation, we compared the candidates on the basis of binding performance and biochemical quality. Pairwise epitope competition among the five clones is summarized in **Supplementary Fig. 7a**. SEC-HPLC showed monomeric contents of 91.8%, 68.7%, 85.1%, 80.7%, and 93.9% for CLCM01, CLCM02, CLCM04, CLCM05, and CLCM08, respectively (**Supplementary Fig. 7b**). CLCM05 and CLCM08 had similar recombinant-antigen binding potency, and CLCM08 carried 13.2 percentage points higher monomeric content together with a favorable binding-to-display profile. We therefore selected CLCM08 as the anti-CD222 targeting antibody for LNP conjugation (**Fig. 1h**).

### 3.2 Engineering and physicochemical characterization of antibody-conjugated mRNA-LNPs

CLCM08 and anti-CD5 antibodies were functionalized with DBCO and coupled to azide-bearing mRNA-LNPs by strain-promoted azide-alkyne cycloaddition (**Fig. 2a**) [49]. Untargeted, anti-CD222, anti-CD5, and dual anti-CD222/anti-CD5 formulations were generated on a common SM-102-based lipid core [47], so that all downstream comparisons differ in targeting ligand rather than in lipid composition.

**Fig. 2.**
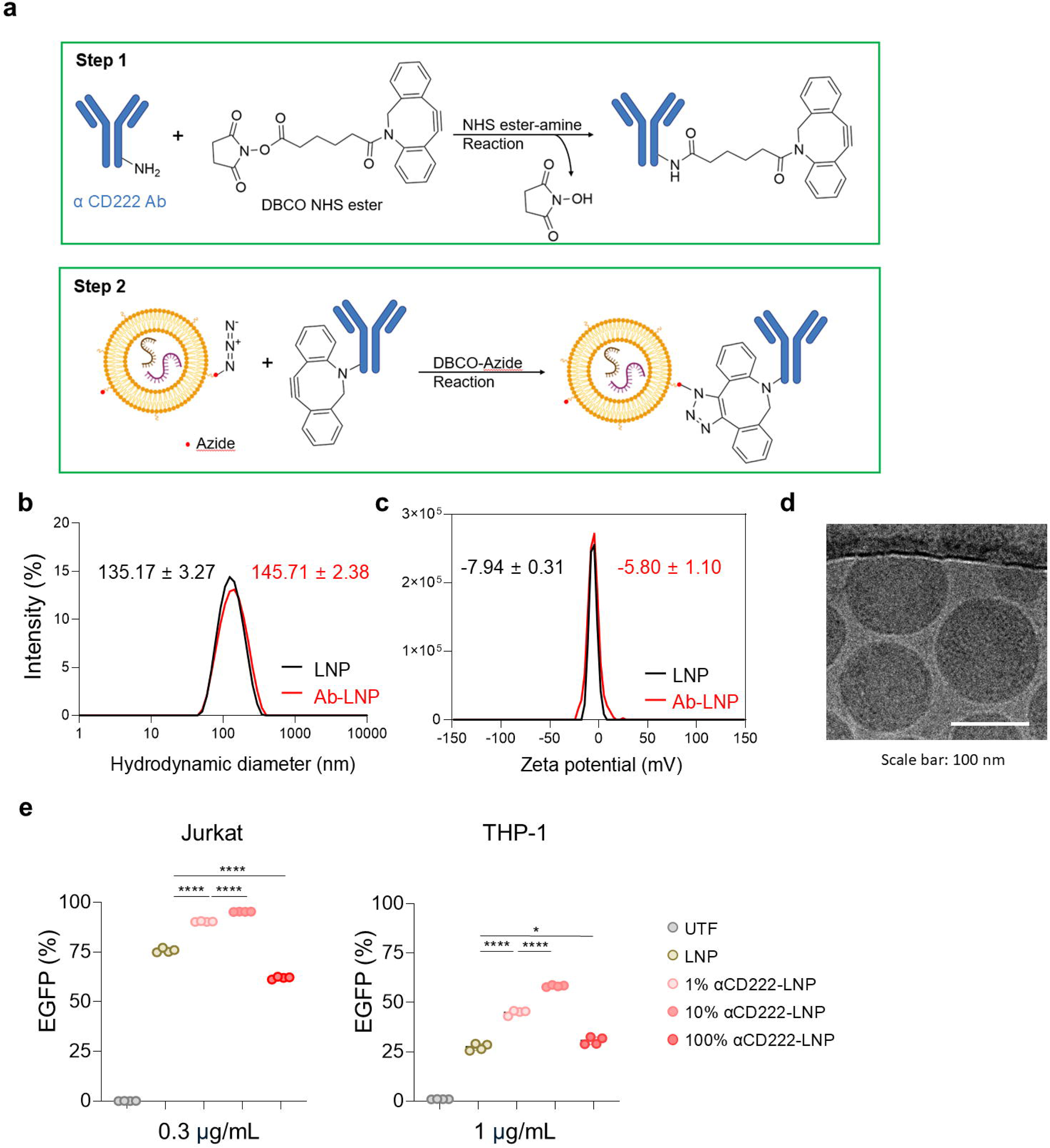
Engineering and physicochemical characterization of antibody-conjugated mRNA-LNPs. (a) Two- step conjugation scheme: antibodies are functionalized with DBCO and coupled to azide-bearing mRNA- LNPs by strain-promoted azide-alkyne cycloaddition. (b) Hydrodynamic size distributions of untargeted and anti-CD222/anti-CD5 antibody-conjugated LNPs by dynamic light scattering. (c) Zeta potential distributions. (d) Cryo-transmission electron micrograph of antibody-conjugated LNPs. Scale bar, 100 nm. (e) EGFP mRNA delivery as a function of nominal anti-CD222 antibody input (1%, 10% and 100%) in Jurkat cells at 0.3 μg/mL and THP-1 cells at 1 μg/mL. Numerical values for size, polydispersity index, zeta potential and encapsulation efficiency of every formulation are given in Supplementary Table S3. Data are presented as mean ± SD from n = 4 independent experiments. Statistical significance for the data in panel (e) was determined by one-way ANOVA followed by Tukey’s multiple-comparison test. ns, not significant; *P < 0.05; **P < 0.01; ***P < 0.001; ****P < 0.0001.

Linker identity was evaluated as an independent formulation variable (**Supplementary Fig. 8a–c**). In the reporter screen, the shortest C2 spacer produced the highest signal, approximately 2,200 arbitrary units, whereas the PEG24 condition fell below the reference control used in the screen (**Supplementary Fig. 8c**). Antibody-particle spacing therefore influences delivery, and the C2 spacer was carried forward for every conjugate reported in this study.

Across the four formulations, bulk particle properties were closely matched, so that the formulation comparisons that follow differ in targeting ligand rather than in particle physicochemistry. Antibody conjugation shifted hydrodynamic diameter from 135.17 ± 3.27 nm for untargeted LNP to 145.71 ± 2.38 nm for the dual-targeted formulation, with PDI ranging from 0.126 to 0.194 and zeta potential between - 7.94 and -3.95 mV for every formulation (**Fig. 2b,c** and **Supplementary Table S3**). Encapsulation efficiency was 75% or greater in all cases. Complete values are tabulated in **Supplementary Table S3**. Cryo-TEM showed discrete particles with the morphology expected for this lipid composition (**Fig. 2d**).

CLCM08 input was then titrated at nominal ratios of 1%, 10%, and 100% (**Fig. 2e**). At the 10% input, the EGFP-positive fraction reached approximately 95% in Jurkat cells at 0.3 μg/mL mRNA and 58% in THP- 1 cells at 1 μg/mL. At the 100% input, delivery fell to approximately 62% and 30%, respectively, approaching or falling below the untargeted-LNP condition. Because post-purification antibody molecules per particle were not quantified, these conditions are described throughout as nominal antibody-input ratios rather than as measured surface densities. On that basis, delivery was non-monotonic with antibody input, and the 10% condition was carried forward for all subsequent experiments.

### 3.3 Targeting-ligand ranking inverts between PBMC monocytes and isolated activated T cells

Figure 3a schematically summarizes the four EGFP mRNA-loaded LNP formulations evaluated in this study: untargeted LNP, anti-CD222 LNP, anti-CD5 LNP, and dual anti-CD222/anti-CD5 LNP. The three conjugated formulations were built on the same SM-102 core with the same C2 linker and the same nominal total IgG input, and all four particles were matched in hydrodynamic diameter, polydispersity and zeta potential (**Fig. 2b,c** and **Supplementary Table S3**). The targeting ligand is therefore the only variable that separates them, and any difference in the comparisons that follow reports the ligand- recipient-cell pairing rather than the formulation. Using the conjugation condition selected above (**Fig. 2e**), we asked whether the ranking of these four formulations is stable across recipient-cell preparations.

**Fig. 3.**
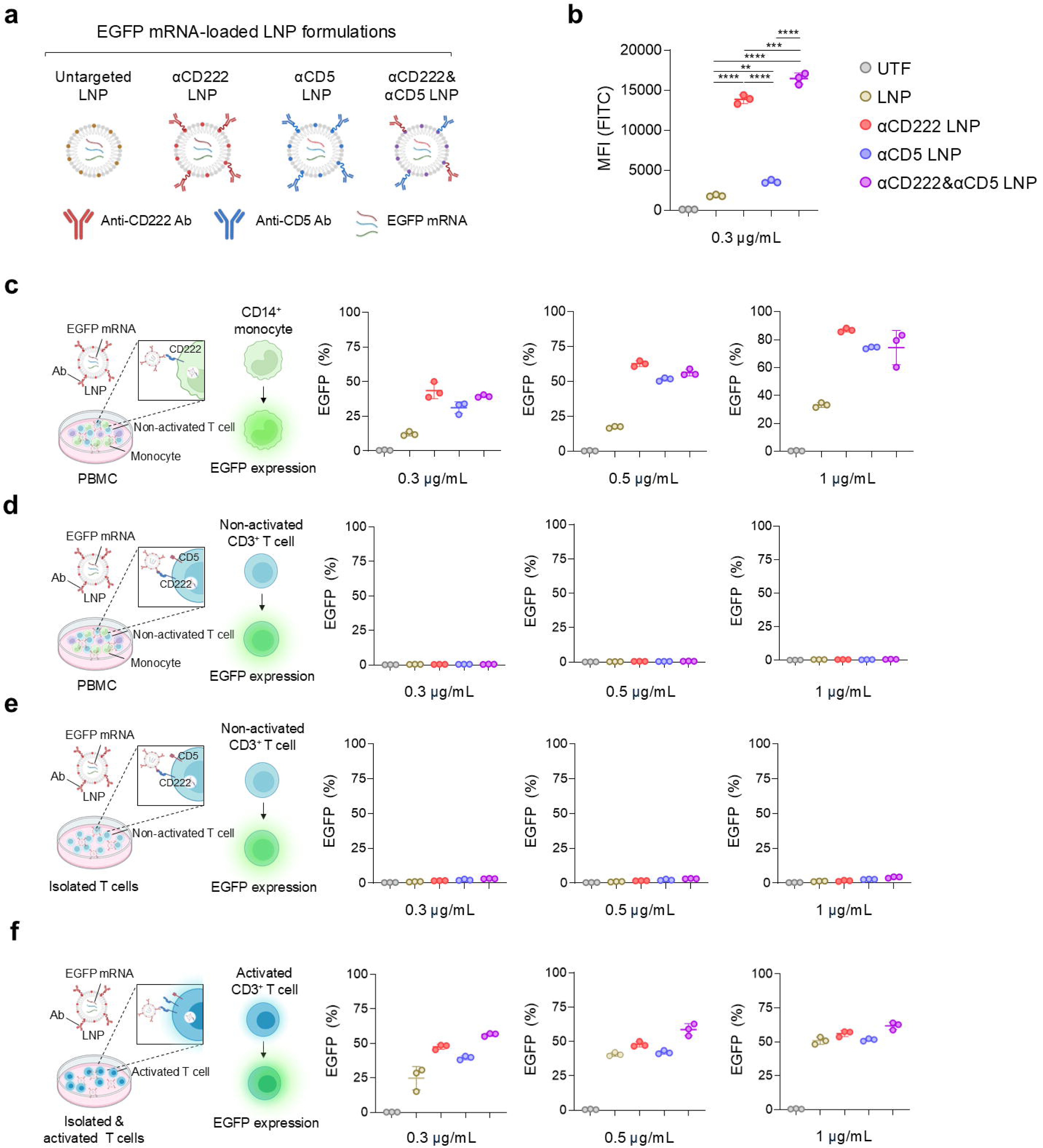
Targeting-ligand ranking inverts between PBMC monocytes and isolated activated T cells. (a) EGFP mRNA-loaded LNP formulations compared in this study. (b) EGFP delivery to Jurkat cells at 0.3 μg/mL. (c) EGFP-positive fraction of CD14-positive monocytes within non-activated PBMC cultures at three mRNA doses. (d) EGFP-positive fraction of CD3-positive T cells within the same PBMC cultures. (e) EGFP-positive fraction of isolated non-activated T cells. (f) EGFP-positive fraction of isolated CD3/CD28-activated T cells. Primary-cell data are presented as mean ± SD of n = 3 technical replicates from a single donor. No inferential statistics were applied (Methods 2.24). Jurkat data (panel b) are presented as mean ± SD from n = 3 independent experiments and were compared by one-way ANOVA followed by Tukey’s multiple-comparison test, and the significance key applies to panel (b) only. ns, not significant; *P < 0.05; **P < 0.01; ***P < 0.001; ****P < 0.0001. UTF, untransfected cells; LNP, untargeted LNP.

In Jurkat cells treated with 0.3 μg/mL mRNA, anti-CD222 LNP, anti-CD5 LNP, and dual anti- CD222/anti-CD5 LNP produced approximately 7.7-, 2.0-, and 9.1-fold higher EGFP MFI than untargeted LNP, respectively (**Fig. 3b**). Untargeted LNP mixed with free anti-CD222 or free anti-CD5 antibody did not reproduce these gains, confirming that covalent particle display, and not the presence of soluble antibody, drives the enhancement (**Supplementary Fig. 9a**). Anti-CD222-containing formulations therefore produced the largest enhancement in this cell line, and anti-CD5 LNP alone produced a 2.0-fold increase.

To further assess the contribution of receptor engagement to targeted delivery, soluble-antibody competition experiments were performed in Jurkat cells. Excess soluble anti-CD222 reduced anti-CD222- LNP delivery by approximately 78%, whereas combined soluble anti-CD222 and anti-CD5 reduced dual- LNP delivery by approximately 77% (**Supplementary Fig. 9b**). In contrast, competition of anti-CD5-LNP delivery produced only a partial reduction, suggesting that CD5 recognition alone may not fully account for the delivery observed with this formulation.

To examine whether the targeting enhancement was observed across different recipient-cell contexts, SNU-5 cells were included as an additional comparator. In Jurkat cells, anti-CD222 and dual anti- CD222/anti-CD5 LNPs markedly increased EGFP signal relative to untargeted LNP, whereas no clear increase was observed among the targeted formulations in SNU-5 cells under the same normalized comparison (**Supplementary Fig. 9c**). Because surface CD222 and CD5 abundance was not quantified in parallel in the current experiment, the SNU-5 comparison is considered supportive, rather than definitive, evidence that the delivery enhancement depends on the targeting ligand–recipient-cell context.

We next investigated whether the enhanced delivery observed in the monocyte compartment was mediated by CD222 recognition or by interactions between the antibody Fc region and Fcγ receptors expressed by monocytes. To separate these contributions, an Fc-silenced LALA-PG variant of CLCM08 was generated and conjugated to EGFP mRNA-LNPs. In CD14-positive monocytes within PBMC cultures, Fc-silenced anti-CD222 LNP increased both the EGFP-positive fraction and EGFP MFI relative to untargeted LNP and to an Fc-silenced isotype-control LNP (**Supplementary Fig. 10a**), establishing a CD222-dependent component of uptake and productive mRNA delivery. Wild-type anti-CD222 LNP nevertheless produced higher reporter expression than the Fc-silenced variant, consistent with an additional Fc-associated contribution in Fcγ receptor-expressing monocytes. Under the tested conditions, CD222 recognition was therefore sufficient to enhance monocyte delivery, and retention of the wild-type Fc added further to that effect.

Soluble-antibody competition was subsequently used to further assess receptor involvement in primary immune-cell preparations. In CD14-positive monocytes within PBMC cultures, excess soluble anti- CD222 antibody reduced anti-CD222-LNP-mediated EGFP expression, and combined soluble anti-CD222 and anti-CD5 antibodies reduced delivery by the dual-targeted formulation. Competition of anti-CD5-LNP delivery was more limited (**Supplementary Fig. 10b**). Similar competition experiments in isolated CD3/CD28-activated T cells showed reduced delivery of anti-CD222, anti-CD5, and dual-targeted LNPs following treatment with the corresponding soluble antibodies (**Supplementary Fig. 10c**). Together, these results support receptor-dependent contributions to targeted delivery while indicating that the magnitude of competition differs according to the targeting ligand and recipient-cell preparation.

Having evaluated targeting in cell-line models, we next examined primary human immune-cell populations. All primary-cell comparisons below are from a single donor and are reported descriptively (Methods 2.24). In PBMC cultures recovered in IL-2 and not exposed to CD3/CD28 stimulation (hereafter non-activated PBMC cultures), CD14-positive monocytes showed dose-dependent EGFP expression (**Fig. 3c** and **Supplementary Fig. 11a**). Anti-CD222 LNP reached approximately 44%, 63%, and 87% EGFP-positive cells at 0.3, 0.5, and 1.0 μg/mL, compared with approximately 12%, 18%, and 33% for untargeted LNP, corresponding to numerical gains of approximately 3.7-, 3.5-, and 2.6-fold in this donor. The dual LNP reached approximately 40%, 57%, and 74%, and anti-CD5 LNP reached approximately 31%, 52%, and 74%. Because monocytes do not provide a canonical CD5-positive target population, the enhanced monocyte delivery observed with anti-CD5-containing particles is not assigned to antigen-specific recognition. The equivalent Fc-variant comparison was performed only for anti-CD222 (**Supplementary Fig. 10a**), so the origin of the anti-CD5-associated monocyte signal is not resolved by the present data.

CD3-positive T cells in the same non-activated PBMC cultures remained near background for EGFP expression across formulations and doses (**Fig. 3d**). To determine whether this limited delivery was attributable to the PBMC environment, T cells were isolated and exposed to the same LNP formulations without CD3/CD28 activation. These isolated non-activated T cells likewise showed minimal EGFP expression (**Fig. 3e**), so isolation alone did not enhance mRNA delivery. Isolated CD3/CD28-activated T cells, in contrast, were transfected substantially more efficiently, and the dual-targeted formulation produced the largest gain (**Fig. 3f** and **Supplementary Fig. 11b**). The dual LNP reached approximately 56%, 59%, and 62% EGFP-positive cells at 0.3, 0.5, and 1.0 μg/mL. By MFI, neither single-ligand LNP differed clearly from untargeted LNP at 0.5 or 1.0 μg/mL, whereas the dual formulation remained higher. The relative gain narrowed with increasing dose, from approximately 2.3-fold over untargeted LNP at 0.3 μg/mL to approximately 1.2-fold at 1.0 μg/mL.

The ranking therefore inverts between the two preparations. Anti-CD222 LNP delivered 2.6- to 3.7-fold more reporter than untargeted LNP in CD14-positive monocytes within non-activated PBMC cultures, and lost that advantage in isolated activated T cells, where by MFI it did not differ clearly from untargeted LNP at 0.5 or 1.0 μg/mL and the dual formulation produced the largest gain. Cell type, isolation and activation state were varied together because these are the formats in which immune cells are actually prepared for ex vivo engineering; the inversion is therefore a property of the recipient-cell preparation as a whole rather than of activation alone.

### 3.4 CAR mRNA delivery reproduces the inversion and reveals a payload-specific deviation

To determine whether the recipient-cell-dependent delivery patterns observed with EGFP extended to a functional CAR payload, CD19 CAR mRNA was encapsulated in the same four LNP formulations, prepared and conjugated under the conditions used for the reporter experiments. The CAR construct contained an extracellular MYC tag, enabling flow-cytometric quantification of surface CAR expression (**Fig. 4a**). In Jurkat cells at 0.3 μg/mL, all transfected conditions exceeded 95% CAR-positive cells, so MFI was used as the principal comparative readout. The dual LNP reached approximately 14,192 MFI, 3.6-fold above untargeted LNP at approximately 3,907 MFI. Anti-CD222 and anti-CD5 LNPs reached approximately 8,141 and 6,014 MFI, respectively (**Fig. 4b**).

**Fig. 4.**
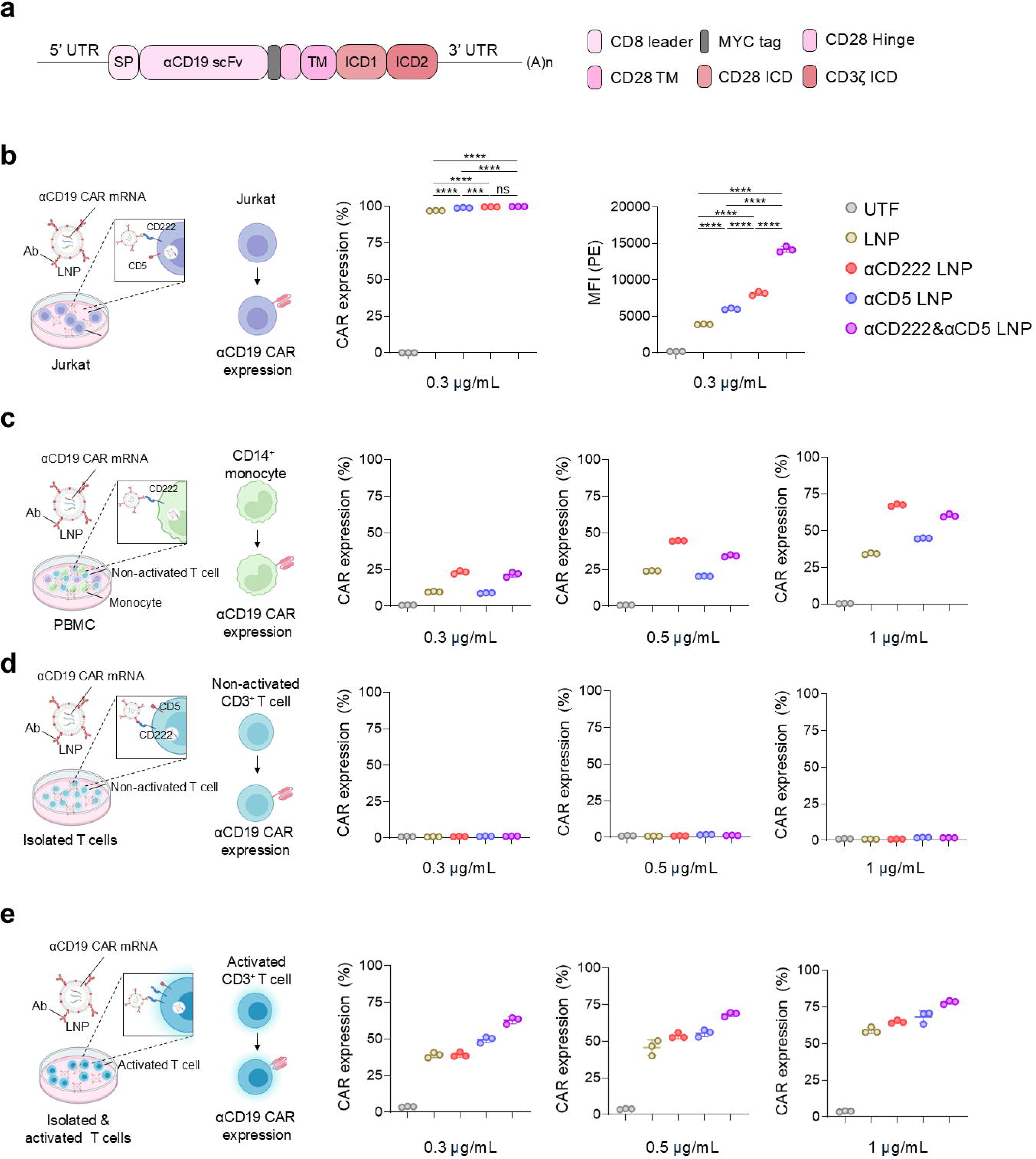
CAR mRNA delivery reproduces the preparation-dependent inversion. (a) Anti-CD19 CAR construct and MYC-tag detection strategy. (b) CAR expression in Jurkat cells at 0.3 μg/mL. (c) CAR expression in CD14-positive monocytes within non-activated PBMC cultures. (d) CAR expression in isolated non-activated CD3-positive T cells. (e) CAR expression in isolated CD3/CD28-activated T cells. Primary-cell data are presented as mean ± SD of n = 3 technical replicates from a single donor. No inferential statistics were applied (Methods 2.24). Jurkat data (panel b) are presented as mean ± SD from n = 3 independent experiments and were compared by one-way ANOVA followed by Tukey’s multiple- comparison test, and the significance key applies to panel (b) only. ns, not significant; *P < 0.05; **P < 0.01; ***P < 0.001; ****P < 0.0001. UTF, untransfected cells; LNP, untargeted LNP.

Having established robust CAR expression in Jurkat cells, we next examined primary immune-cell preparations. In CD14-positive monocytes within non-activated PBMC cultures, anti-CD222 LNP reached approximately 23%, 45%, and 68% CAR-positive cells at 0.3, 0.5, and 1.0 μg/mL (**Fig. 4c** and **Supplementary Fig. 12a**). At 1.0 μg/mL, anti-CD222 LNP reached approximately 68% CAR-positive monocytes, the dual anti-CD222/anti-CD5 LNP approximately 60%, and anti-CD5 LNP approximately 45%, compared with approximately 34% for untargeted LNP. Anti-CD5 LNP did not consistently exceed untargeted LNP across doses in this CAR assay, although an increase was observed at 1.0 μg/mL, in contrast to its reporter-mRNA behavior. This payload-specific deviation is reported explicitly because the EGFP and CAR transcripts differ in length and expression kinetics, and it shows that a formulation ranking established with a reporter does not transfer unchanged to a therapeutic payload.

We next examined whether the T-cell delivery pattern observed with EGFP mRNA was also reflected in the CAR mRNA experiments. Non-activated T cells remained difficult to engineer. Consistent with the reporter results, isolated non-activated T cells showed minimal CAR surface expression across the tested formulations (**Fig. 4d**). These observations are not interpreted as a direct reporter-versus-CAR comparison because the preparations were not matched.

In isolated CD3/CD28-activated T cells, transfection was substantially more efficient, consistent with the EGFP experiments. The dual LNP reached approximately 63%, 68%, and 78% CAR-positive cells at 0.3, 0.5, and 1.0 μg/mL (**Fig. 4e** and **Supplementary Fig. 12b**). At 0.3 μg/mL, anti-CD222 LNP and untargeted LNP were both approximately 39% CAR-positive, while anti-CD5 LNP reached approximately 50% and the dual LNP 63%. At 1.0 μg/mL, the dual LNP reached approximately 78% CAR-positive cells, compared with approximately 69% for anti-CD5 LNP, 65% for anti-CD222 LNP and 59% for untargeted LNP; as with the reporter payload, the relative advantage of the dual formulation narrowed with increasing dose. The CAR payload therefore reproduces the inversion seen with the reporter: anti-CD222 LNP delivered 68% CAR-positive monocytes at 1.0 μg/mL yet did not separate from untargeted LNP in activated T cells at 0.3 μg/mL, and the ordering of the single-ligand formulations differed between the two preparations while the dual formulation remained the strongest condition in activated T cells.

### 3.5 Delivery efficiency and inflammatory output are separable properties of the targeting ligand

To determine whether receptor-directed CAR mRNA delivery altered short-term cell fitness or immune-cell state, we measured viability, surface phenotype, and cytokine output using the workflow in **Fig. 5a**. This analysis was intended to distinguish efficient CAR delivery from formulation-associated toxicity or immune activation. Short-term viability remained high across formulations and doses. Annexin V/PI analysis showed approximately 88–93% viable cells in the monocyte gate and 97–98% in activated T cells, with no formulation causing a clear increase in apoptotic fractions under the tested conditions (**Fig. 5b,c** and **Supplementary Fig. 13**).

**Fig. 5.**
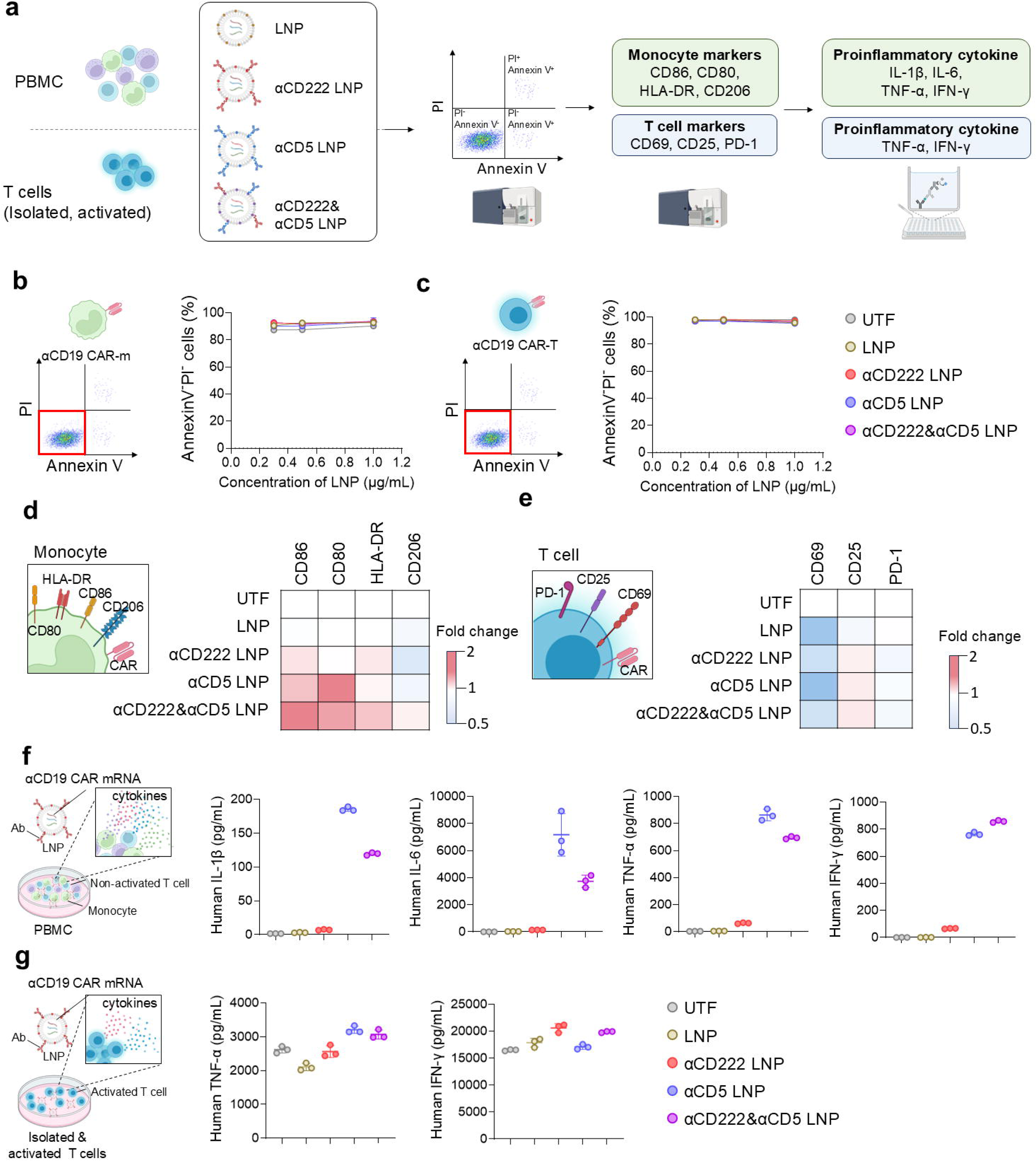
Delivery efficiency and inflammatory output are separable properties of the targeting ligand. (a) Experimental workflow. (b, c) Annexin V/propidium iodide viability of PBMC-derived CD14-positive monocytes (b) and isolated activated T cells (c). (d) Monocyte surface markers CD86, CD80, HLA-DR and CD206. (e) T-cell surface markers CD69, CD25 and PD-1. Data in panels (d) and (e) represent heatmap data expressed as relative values normalized to UTF (set to 1). (f) Cytokine concentrations in non-activated PBMC culture supernatants. (g) Cytokine concentrations in isolated activated T-cell cultures. All panels show primary-cell data. Primary-cell data are presented as mean ± SD of n = 3 technical replicates from a single donor. No inferential statistics were applied (Methods 2.24). UTF, untransfected cells; LNP, untargeted LNP.

We next examined whether LNP treatment altered immune-cell surface phenotype. In non-activated PBMC cultures, monocyte phenotype markers changed modestly with formulation (**Fig. 5d** and **Supplementary Fig. 14a**). The dual LNP increased CD86 by approximately 1.8-fold, CD80 by approximately 1.6-fold, and HLA-DR by approximately 1.4-fold relative to UTF, whereas CD206 remained within a narrow range. These changes were consistent with a shift toward an activated antigen-presenting phenotype but were not accompanied by a substantial loss of viability.

In isolated activated T cells, CD25, CD69, and PD-1 showed only small formulation-associated shifts (**Fig. 5e** and **Supplementary Fig. 14b**). CD25 increased modestly, CD69 was lower than in untreated cells across LNP conditions, and PD-1 remained essentially unchanged. Given the magnitude of these differences relative to the assay range, they are not interpreted as evidence of a major change in T-cell activation state.

Because antibody targeting may alter inflammatory output independently of delivery efficiency, we measured cytokines released after CAR mRNA-LNP treatment. In mixed PBMC cultures, cytokine output diverged sharply with targeting-ligand composition (**Fig. 5f**). Relative to untargeted LNP, anti-CD5 LNP raised IL-6 from approximately 31 to 7,178 pg/mL, IL-1β from approximately 3 to 186 pg/mL, and TNF-α from approximately 5 to 864 pg/mL, corresponding to increases of approximately 233-, 59-, and 167-fold. Because the untargeted-LNP baseline concentrations were low, these ratios are reported together with the absolute concentrations. IFN-γ reached approximately 760 pg/mL with anti-CD5 LNP and 850 pg/mL with the dual LNP, compared with approximately 20 pg/mL for untargeted LNP. Anti-CD222 LNP reached an IL-6 concentration of approximately 139 pg/mL, 4.5-fold over untargeted LNP and 52-fold below anti-CD5 LNP, and the dual formulation produced intermediate values. On one lipid core and at matched nominal IgG input, the choice of targeting ligand therefore changed IL-6 output by approximately 52-fold between the two single-ligand formulations.

The pronounced cytokine response was observed in mixed PBMC cultures containing monocytes and was markedly attenuated in isolated T-cell cultures. In the latter setting, anti-CD5 LNP raised TNF-α only approximately 1.5-fold, from approximately 2,108 to 3,208 pg/mL, and did not materially change IFN-γ (**Fig. 5g**). The present study was not designed to distinguish Fcγ receptor-dependent particle recognition from CD5-dependent signaling or other preparation-associated effects, and the cytokine data are interpreted as a formulation-level biological output rather than as evidence for a specific receptor-signaling pathway. The pronounced response was therefore observed only in the myeloid-containing culture; because IL-6 and IL-1β were not measured in the isolated T-cell condition, this myeloid dependence is inferred from the culture composition rather than measured directly. The Fc-variant comparison in Supplementary Fig. 10a addresses the receptor-level question for anti-CD222 only.

### 3.6 Effector activity after CAR mRNA delivery

To determine whether CAR mRNA delivery to the PBMC-derived monocyte compartment translated into target-cell uptake, CAR mRNA-LNP-treated PBMCs were co-cultured with pHrodo Red-labeled CD19- positive Raji cells and pHrodo-positive events were quantified within CD14-positive monocytes (**Fig. 6a,b**). Anti-CD222-LNP-treated cultures reached approximately 24%, 37%, and 63% pHrodo-positive monocytes at nominal effector-to-target ratios of 10:1, 5:1, and 1:1, respectively. Untransfected and untargeted-LNP background rose in parallel with target input, reaching approximately 39% in untransfected cultures at the nominal 1:1 condition, so the lower-background 10:1 condition provides the most interpretable comparison. At the 10:1 condition, all three antibody-conjugated formulations exceeded untargeted LNP: anti-CD222 LNP produced approximately 23.8% pHrodo-positive monocytes, the dual anti-CD222/anti-CD5 LNP approximately 21%, and anti-CD5 LNP approximately 17%, compared with approximately 14.8% for untargeted LNP and 4.8% for untransfected cultures. Anti-CD222 LNP therefore produced the largest increase, approximately 1.6-fold over untargeted LNP, and the ordering of the formulations in the monocyte compartment matched the ordering observed for CAR delivery in the same compartment (Fig. 4c), with the dual formulation intermediate. Cytokine profiles after Raji coculture again showed greater inflammatory responses with anti-CD5-containing formulations than with anti- CD222 LNP (**Fig. 6c**). Anti-CD5 LNP produced the highest IL-1β and IL-6 levels, whereas the dual anti- CD222/anti-CD5 LNP produced the highest TNF-α and IFN-γ levels.

**Fig. 6.**
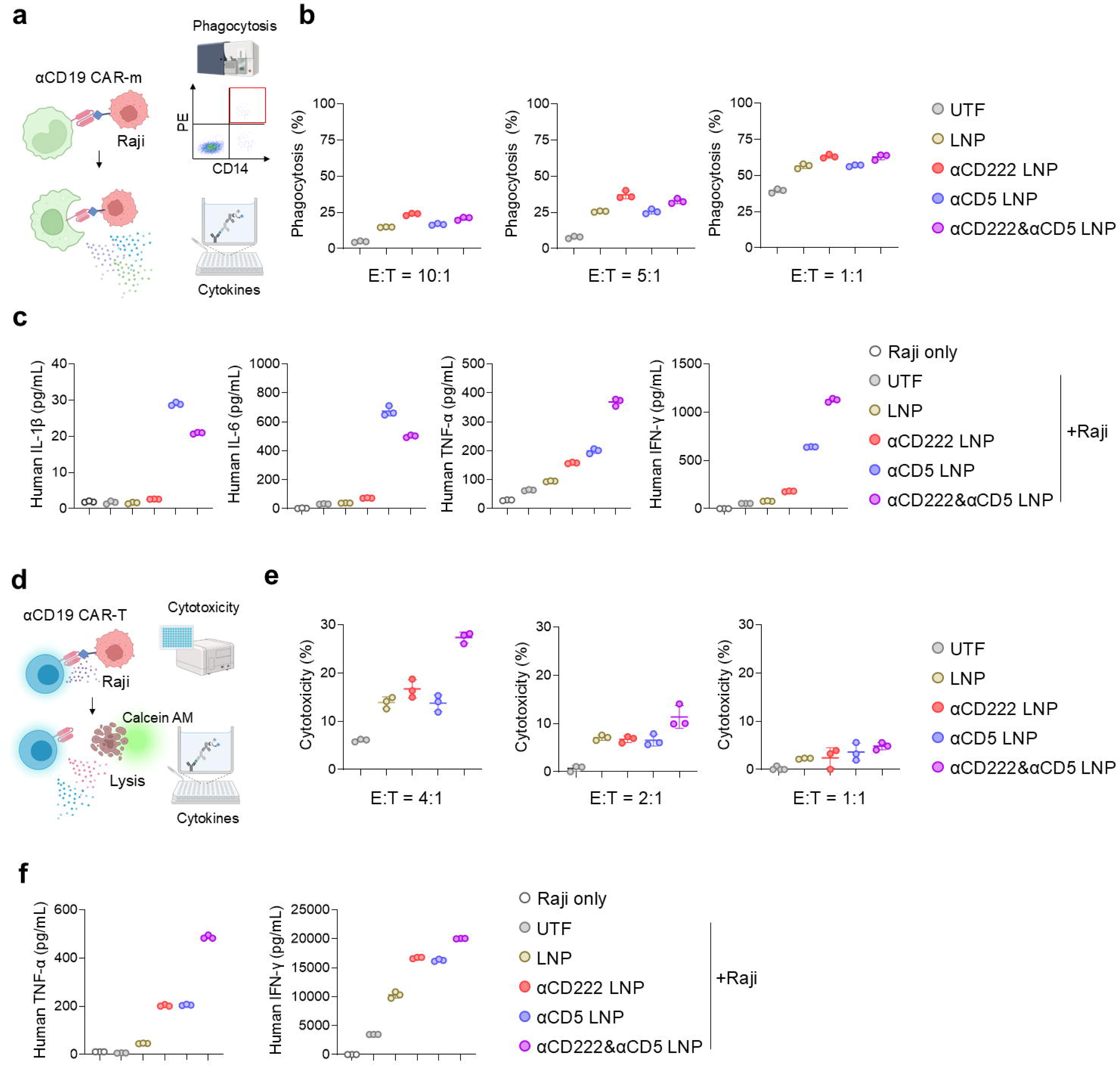
Effector activity after CAR mRNA delivery to monocyte and T-cell preparations. (a) Phagocytosis assay schematic. (b) pHrodo-positive fraction of CD14-positive monocytes after coculture with CD19- positive Raji cells at three nominal effector-to-target ratios, shown with untransfected and untargeted-LNP controls at each ratio. (c) Cytokine concentrations after coculture. (d) Cytotoxicity assay schematic. (e) Calculated specific lysis of calcein-labeled Raji cells by CAR mRNA-LNP-treated activated T cells. (f) Cytokine concentrations after coculture. All panels show primary-cell data. Primary-cell data are presented as mean ± SD of n = 3 technical replicates from a single donor. No inferential statistics were applied (Methods 2.24). UTF, untransfected cells; LNP, untargeted LNP.

We next assessed the functional activity of CAR-engineered activated primary T cells against Raji target cells using a calcein-release cytotoxicity assay (**Fig. 6d,e**). The dual formulation produced approximately 27.4%, 11.4%, and 4.8% calculated specific lysis at effector-to-target ratios of 4:1, 2:1, and 1:1, respectively. At 4:1, this was approximately twofold above untargeted LNP. Neither single-ligand formulation separated clearly from untargeted LNP at any tested ratio — anti-CD222 LNP reached approximately 17% specific lysis versus 14% for untargeted LNP at 4:1, within the spread of technical replicates — and the dual formulation also produced the largest post-coculture TNF-α and IFN-γ increases (**Fig. 6f**). Because the dual formulation generated the largest CAR-positive fraction before coculture, these functional differences are not interpreted independently of CAR-positive effector abundance.

Because antigen-negative target cells were not included, the antigen-specific component of this activity is not quantified here. Within that scope, CAR mRNA delivered from a single LNP platform produced effector activity in both compartments: CAR-engineered monocytes took up CD19-positive Raji cells and CAR-engineered T cells lysed them.

## 4. Discussion

This study shows that the ranking of antibody-targeted mRNA-LNPs inverts with the recipient-cell preparation. A single targeting optimum did not transfer across the immune-cell preparations tested on a common lipid core. Anti-CD222 LNP raised reporter delivery 2.6- to 3.7-fold over untargeted LNP in CD14-positive monocytes within non-activated PBMC cultures, and reached 68% CAR-positive monocytes at 1.0 μg/mL. The same particle matched untargeted LNP in isolated CD3/CD28-activated T cells, both reaching approximately 39% CAR-positive cells at 0.3 μg/mL, where the dual anti-CD222/anti- CD5 formulation instead produced the largest gain. Targeting performance is therefore a property of the ligand-recipient-cell pair rather than an intrinsic property of the targeting antibody, and a ligand ranking measured in one preparation cannot be carried into another without re-testing [20,21,50].

Receptor accessibility and trafficking provide a mechanistic frame for this inversion. CD222/CI-MPR is broadly expressed and constitutively traffics through endosomal compartments [26,27,30], so it supplies endocytic flux wherever it is accessible on the cell surface. Primary monocytes carried the largest accessible CD222 pool in the antibody-binding experiments and showed the largest anti-CD222 delivery gain. Non-activated T cells showed little accessible CD222 and were poorly transfected by every formulation. CD3/CD28 activation raised the CD222-positive T-cell fraction from 8.73% to 92.3% in a representative sample, and in that preparation the dual formulation performed best. Receptor accessibility, set by how the recipient cell is prepared ex vivo, therefore tracks the inversion. Because activation, isolation, and cell type were varied together, and because the primary-cell data derive from one donor, these observations support a preparation-context model rather than a demonstrated general mechanism.

Productive mRNA delivery requires both cell entry and endosomal escape [31]. We did not directly measure intracellular trafficking or cytosolic escape, so we do not assign a receptor-specific escape mechanism. Two formulation observations nevertheless emphasize that entry alone is insufficient. First, reporter delivery was non-monotonic with antibody input and peaked at the 10% nominal input rather than the 100% condition. Second, linker identity altered reporter output, with the shortest C2 spacer outperforming longer linkers in the initial screen. That screen was performed with a model IgG in RAW 264.7 cells rather than with the anti-CD222 or anti-CD5 conjugates themselves, so the selected C2 spacer was not re-optimized for the targeting antibodies and recipient-cell preparations used here. This result is consistent with the principle that interfacial design can influence cellular interactions and delivery behavior [17,51–53]. These variables should therefore be treated as independent engineering parameters that can alter the conversion of receptor engagement into productive cytosolic delivery.

Delivery efficiency and inflammatory cost were separable. On one lipid core and at matched nominal IgG input, anti-CD5 LNP raised PBMC IL-6 to 7,178 pg/mL while anti-CD222 LNP reached 139 pg/mL, a 52-fold difference, even though the two formulations differed by 13 percentage points in monocyte reporter delivery at 1.0 μg/mL (87% versus 74%). The same anti-CD5 formulation raised TNF-α only 1.5-fold in isolated T-cell cultures. IL-6 and IL-1β were not measured in the isolated T-cell condition, so the myeloid dependence of the IL-6 divergence is inferred from the culture composition rather than measured directly [54–56]. All PBMC cultures were recovered in IL-2 before treatment, so this comparison is made on a common cytokine background rather than against a truly resting culture. Whatever the underlying mechanism, the magnitude of the divergence establishes that delivery efficiency alone is an insufficient basis for formulation selection, and that inflammatory output must be measured as a second, independent axis during ligand screening.

Part of the monocyte gain is Fc-associated rather than antigen-directed, and this bounds the CD222- specific claim. Fc-silenced anti-CD222 LNP retained a delivery advantage over untargeted LNP and over an Fc-silenced isotype control, which establishes a CD222-dependent component. Wild-type anti-CD222 LNP nevertheless outperformed the Fc-silenced variant, and anti-CD5 LNP raised monocyte reporter delivery to 31–74% in a population that does not express CD5. The most parsimonious reading is that particle-displayed wild-type IgG contributes an antigen-independent Fcγ receptor-mediated component in monocytes [57], on top of the antigen-directed component. Completing the wild-type versus LALA-PG [58] comparison for both targeting antibodies, together with a wild-type non-binding isotype control, resolves the two contributions quantitatively and is required before the monocyte gain is attributed to CD222 alone.

Our study differs from prior antibody-targeted mRNA-LNP work in three ways. First, the targeting antibodies were attached covalently through DBCO–azide chemistry, and linker identity and nominal antibody input were treated as explicit formulation variables rather than as fixed conditions [18–20]. Second, CD222 and CD5 were compared head-to-head on a common LNP core rather than optimized in separate systems, so the comparison reports the ligand rather than the formulation. Third, the same four particles were applied across three matched recipient-cell preparations, which is what exposes the inversion; a study that fixes the preparation cannot detect it. This design does not replace in vivo validation, but it is what allows a ligand ranking to be read as a property of the ligand-recipient-cell pair rather than of the particle, and it defines those pairing rules before systemic delivery is attempted.

The CAR payload preserved the inversion but did not preserve every ranking detail. In monocytes, anti- CD5 LNP enhanced EGFP delivery yet did not consistently exceed untargeted LNP for CAR surface expression. Transcript length, translation kinetics and receptor engagement could all contribute [59]. This argues against assuming that a reporter-optimized formulation retains its exact ranking for a therapeutic mRNA, and supports direct optimization with the intended payload.

Three limitations bound the current conclusions. First, all primary-cell comparisons were performed in a single healthy donor with technical replicates, so no inferential statistics were applied and the inversion should be read as a proof-of-concept observation in one donor rather than as a population-level rule; replication across independent donors is the single most informative experiment remaining. Second, antibody comparisons were based on nominal IgG input, because post-purification surface ligand density was not quantitatively resolved; single- and dual-targeted formulations were matched on nominal total IgG input rather than on measured antibody molecules per particle. Third, the effector-function assays used CD19-positive Raji cells without an antigen-negative counterpart, so the antigen-specific component of the observed phagocytosis and cytotoxicity is not quantified, and the study does not establish the molecular mechanism underlying the anti-CD5-associated cytokine response.

Within these boundaries, the study supports a practical design principle for nonviral ex vivo immune-cell engineering: antibody identity, receptor accessibility, cell preparation, linker architecture and antibody loading form one coupled system and must be optimized together. Anti-CD222 targeting delivered efficiently to monocytes at an IL-6 output of 139 pg/mL, whereas the anti-CD5-containing formulations that were required in activated T-cell preparations raised IL-6 to 7,178 pg/mL for anti-CD5 LNP alone, with the dual formulation intermediate. Optimizing delivery efficiency without simultaneously measuring the inflammatory cost of receptor targeting will select a formulation that is efficient in one cellular context and undesirable in another.

## 5. Conclusions

Recipient-cell preparation, not targeting-ligand identity alone, determined both the efficiency and the inflammatory output of antibody-conjugated mRNA-LNP delivery. Anti-CD222 targeting raised reporter delivery 2.6- to 3.7-fold over untargeted LNP in CD14-positive monocytes within non-activated PBMC cultures and raised IL-6 4.5-fold, to 139 pg/mL. In isolated CD3/CD28-activated T cells the same particle matched untargeted LNP and the dual anti-CD222/anti-CD5 formulation instead produced the largest gain, reaching 78% CAR-positive cells at 1.0 μg/mL, while anti-CD5 LNP raised IL-6 to 7,178 pg/mL, 52-fold above anti-CD222 LNP. Ligand ranking therefore inverts with the recipient-cell preparation, and antibody identity, linker architecture, antibody loading and cell preparation should be optimized as one coupled system for nonviral ex vivo immune-cell engineering.

## CRediT authorship contribution statement

Jisu Hong: Investigation, Methodology, Formal analysis, Writing - original draft. Jinwoong Lim: Investigation, Methodology, Formal analysis, Writing - original draft. Minho Park: Investigation, Methodology, Formal analysis. Jimin Song: Investigation. Byung Joon Lee: Investigation. Miyoung Park: Writing, review, editing. Min-Kyoo Shin^:^ Writing, review, editing. Hyungseok Seo: Writing, review, editing. Hyoung Jin Kang: Conceptualization, Supervision, Funding acquisition, Writing - review and editing. Byung-Soo Kim: Conceptualization, Supervision, Funding acquisition, Writing - review and editing. Chang-Han Lee: Conceptualization, Supervision, Funding acquisition, Writing - review and editing.

## Acknowledgements

This work was supported in part by the Bio&Medical Technology Development Program of the National Research Foundation (NRF), funded by the Ministry of Science and ICT (MSIT) of the Korean government (No. RS-2024-00440679 to H.J.K.), the Innovative New Drug Development Program funded by MSIT (No. RS-2026-25527526 to C.-H.L.), the Mid-Career Researcher Program funded by MSIT (No. RS-2023-00278980 to C.-H.L. and RS-2024-00354235 to B.S.K.), and Korean Fund for Regenerative Medicine funded by MSIT/Ministry of Health and Welfare (RS-2026-25501001 to B.S.K), Republic of Korea.

## Data availability

All public single-cell RNA-sequencing datasets analyzed in this study are listed with their accessions in Supplementary Tables S1 and S2. Analysis code is available from the corresponding author upon reasonable request. Sequences of CLCM08 are subject to patent application KR10-2025-0137979 and are available from the corresponding author under a material transfer agreement. All other data are available from the corresponding author upon reasonable request.

## Declaration of generative AI and AI-assisted technologies in the manuscript preparation process

During the preparation of this work, the authors used OpenAI Codex to assist with language editing, structural organization, consistency checking, and reference verification. The authors reviewed and edited all outputs and take full responsibility for the content of the publication.

---

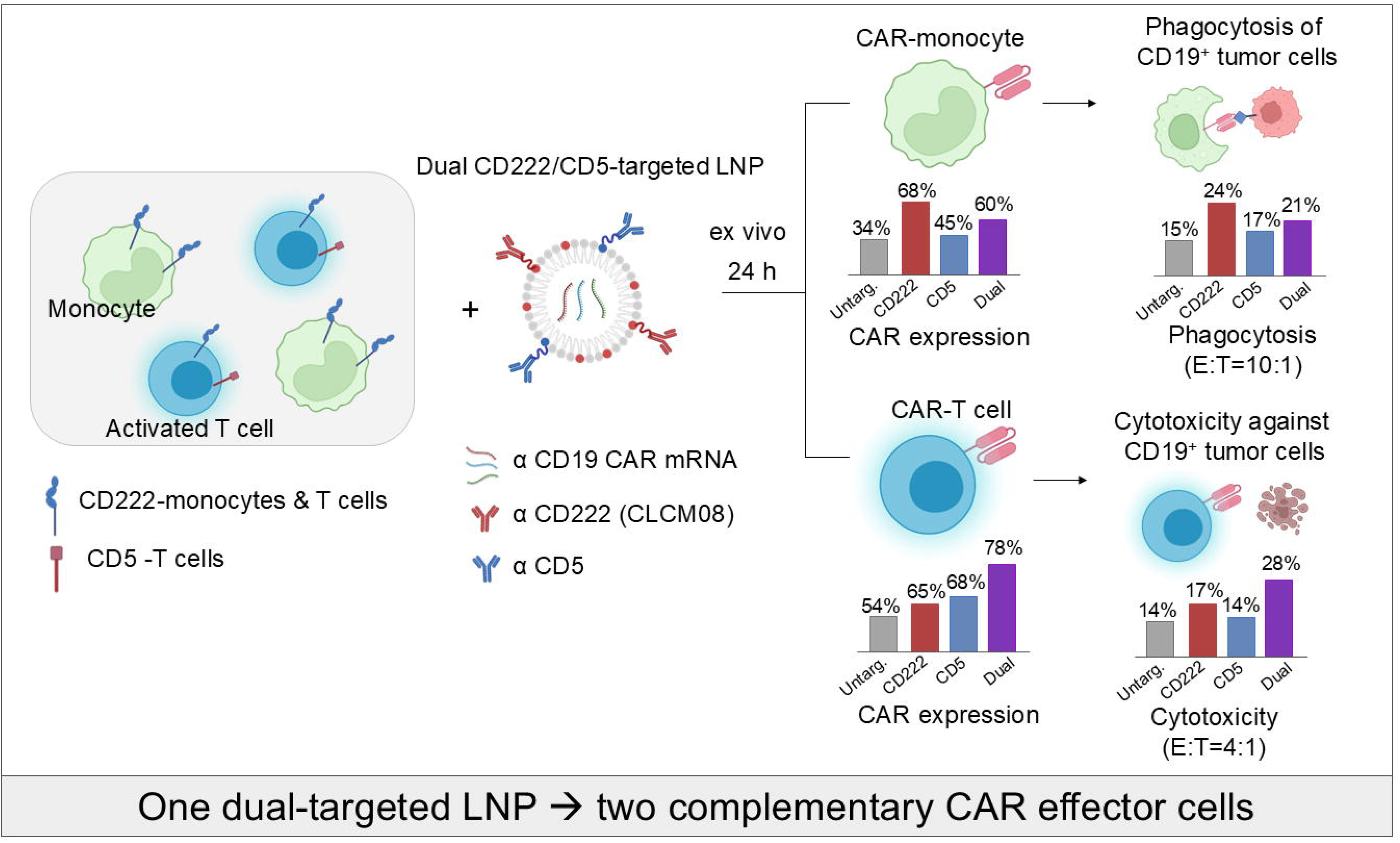

