## Supplementary information for "Dual-targeted nanoparticles enable the ex vivo generation of cytotoxic and phagocytic CAR effector cells"

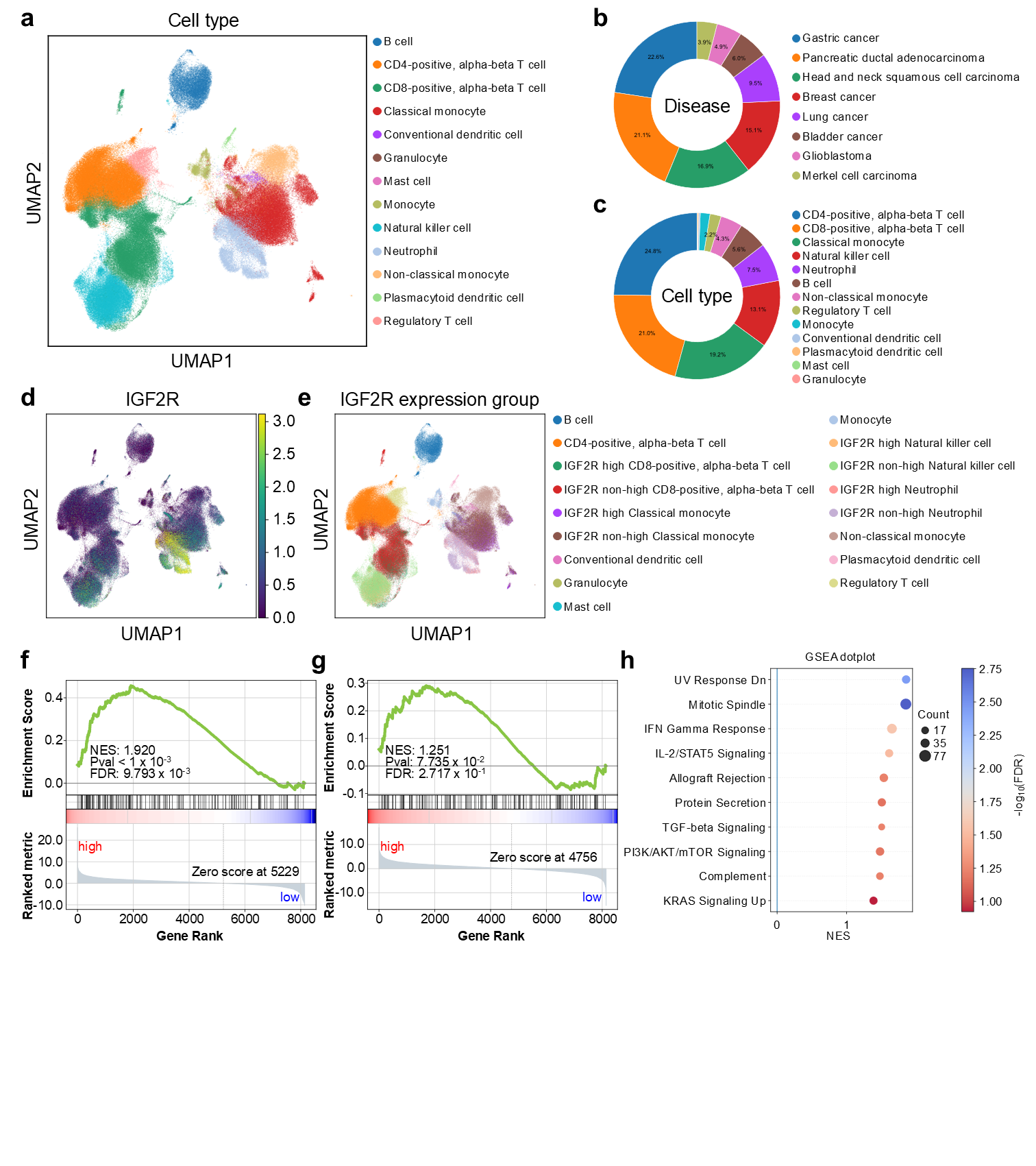

**Supplementary Figure 1. Pan-cancer PBMC single-cell atlas and IGF2R-associated transcriptional programs.**

(a) Harmony-integrated UMAP of 248,266 peripheral blood-derived cells from 77 samples across nine publicly available single-cell RNA-sequencing datasets, annotated into the indicated immune-cell populations. (b) Relative cell-type composition of the integrated atlas. (c) Distribution of cells according to disease origin. (d) UMAP feature plot showing log-normalized IGF2R expression. (e) UMAP showing IGF2R-high and IGF2R-non-high populations; within each cell type, IGF2R-high cells were defined as the top 20% of cells ranked by IGF2R expression. (f) Preranked gene set enrichment analysis (GSEA) of the ENDOCYTOSIS gene set from the KEGG 2026 collection in CD8-positive alpha-beta T cells (NES = 1.920; FDR q = 0.0098). (g) Preranked GSEA of the ENDOCYTOSIS gene set from the KEGG 2026 collection in classical monocytes, showing positive but non-significant enrichment (NES = 1.251; FDR q = 0.2717). (h) Dot plot summarizing the top 10 positively enriched gene sets, ranked by normalized enrichment score (NES), from preranked GSEA using the MSigDB Hallmark 2020 collection in CD8-positive alpha-beta T cells. Dot size represents the number of genes contributing to each gene set, and color indicates −log10(FDR). For GSEA, all tested genes were ranked using the DESeq2 Wald statistic from donor-level pseudobulk comparisons of IGF2R-high and donor-matched IGF2R-non-high populations.

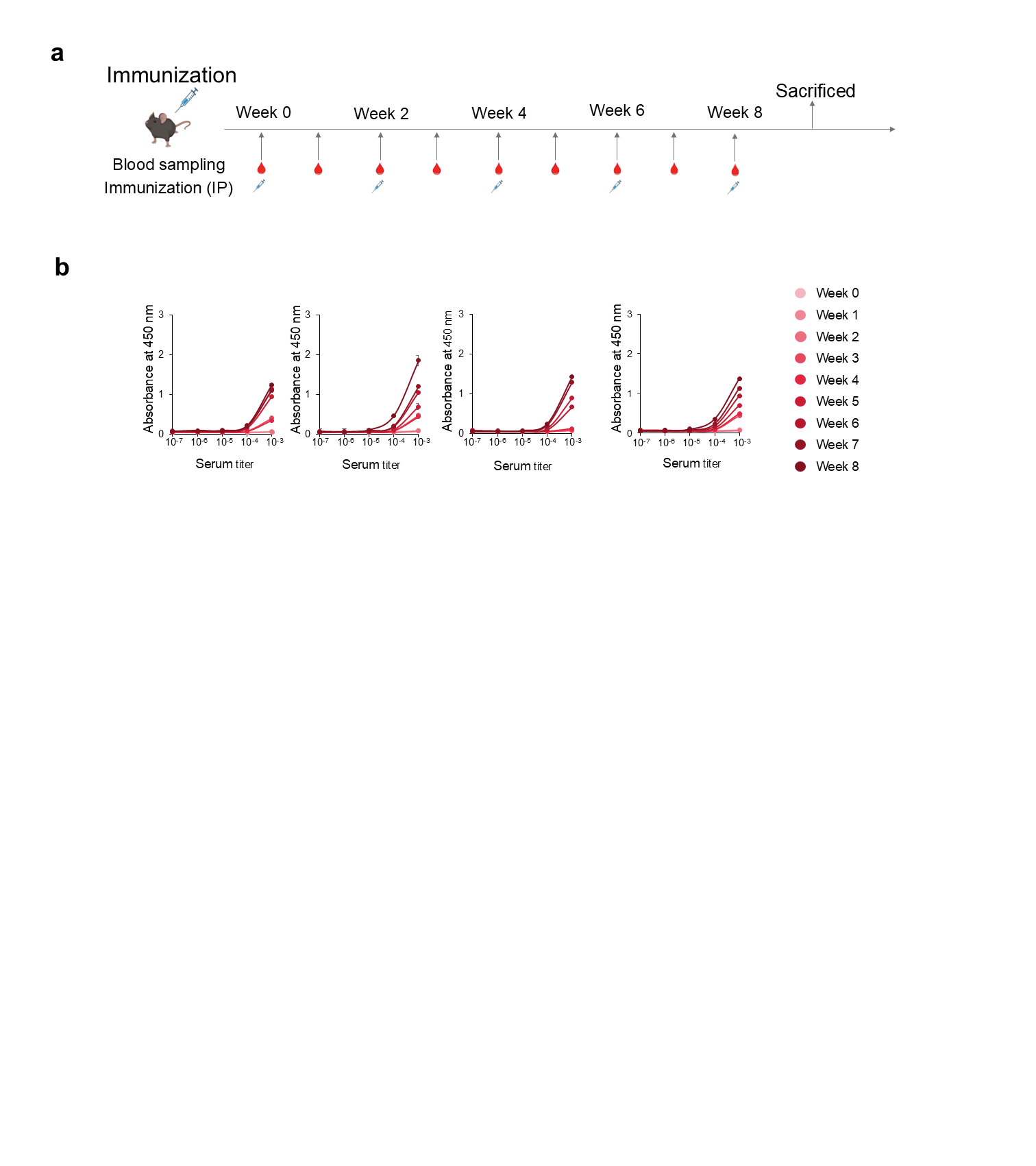

**Supplementary Figure 2. Mouse immunization schedule and longitudinal serum anti-CD222 titers.**

(a) Schematic of the immunization and blood-sampling schedule. Four six-week-old female C57BL/6 mice were immunized intraperitoneally with recombinant human CD222-His formulated with Alum Adjuvant at weeks 0, 2, 4, 6, and 8, and blood was collected weekly. Mice were sacrificed after the final immunization. (b) Longitudinal serum-titer ELISA for the four individual mice. Serially diluted sera collected at weeks 0–8 were tested for binding to immobilized recombinant human CD222-His, and absorbance was measured at 450 nm. Data are presented as mean ± SD of duplicate measurements.

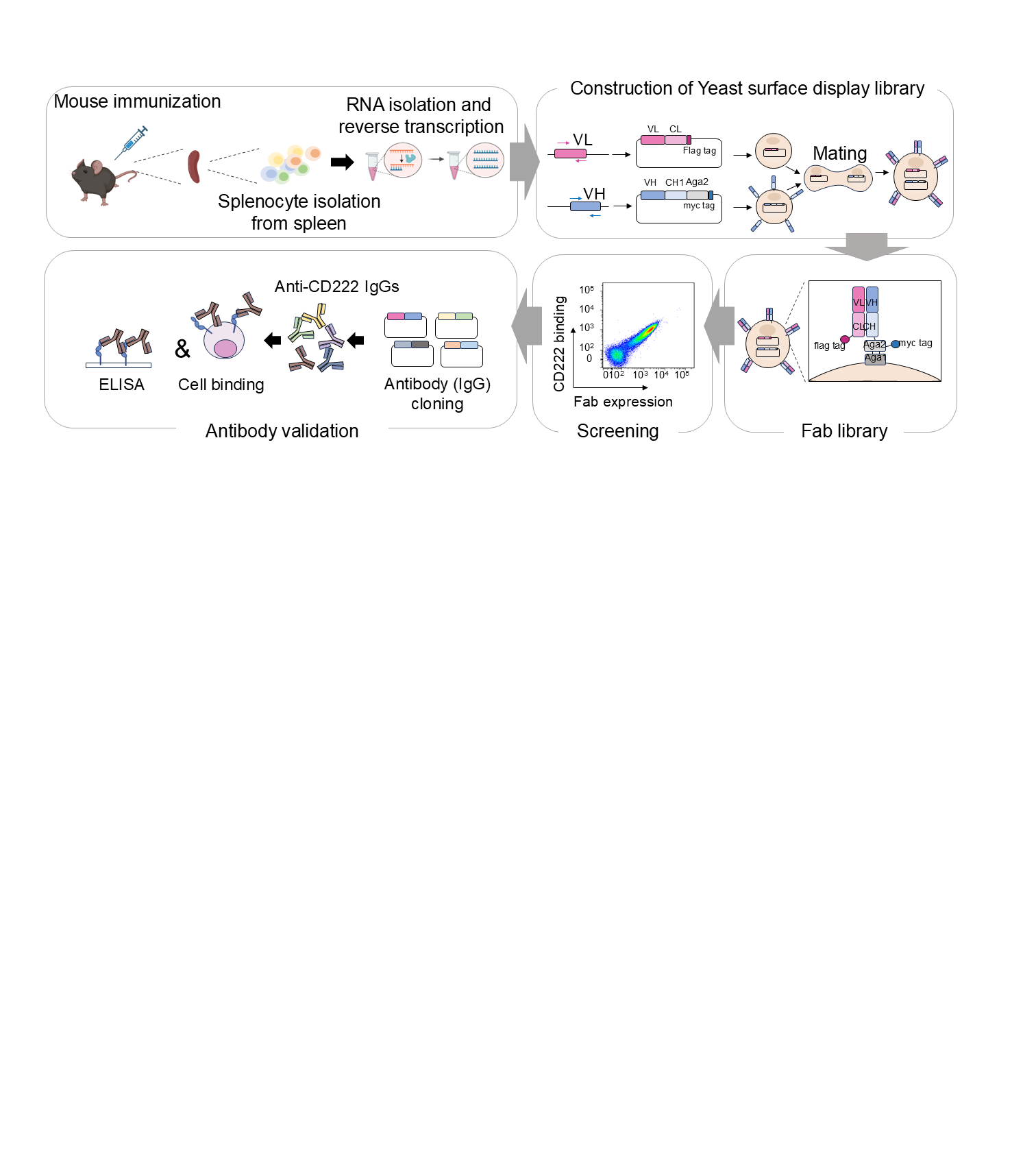

**Supplementary Figure 3. Workflow for immune Fab yeast-display discovery of anti-CD222 antibodies.**

Schematic overview of the anti-CD222 antibody-discovery workflow. Mice were immunized with recombinant human CD222, followed by splenocyte isolation, RNA extraction, and cDNA synthesis. VH and VL repertoires were separately cloned into yeast surface-display vectors and paired by yeast mating to generate an immune Fab library. CD222-binding Fab-expressing yeast cells were enriched by sequential screening, and selected clones were reformatted as human IgG1/κ chimeric antibodies for recombinant-protein ELISA and cell-surface binding validation.

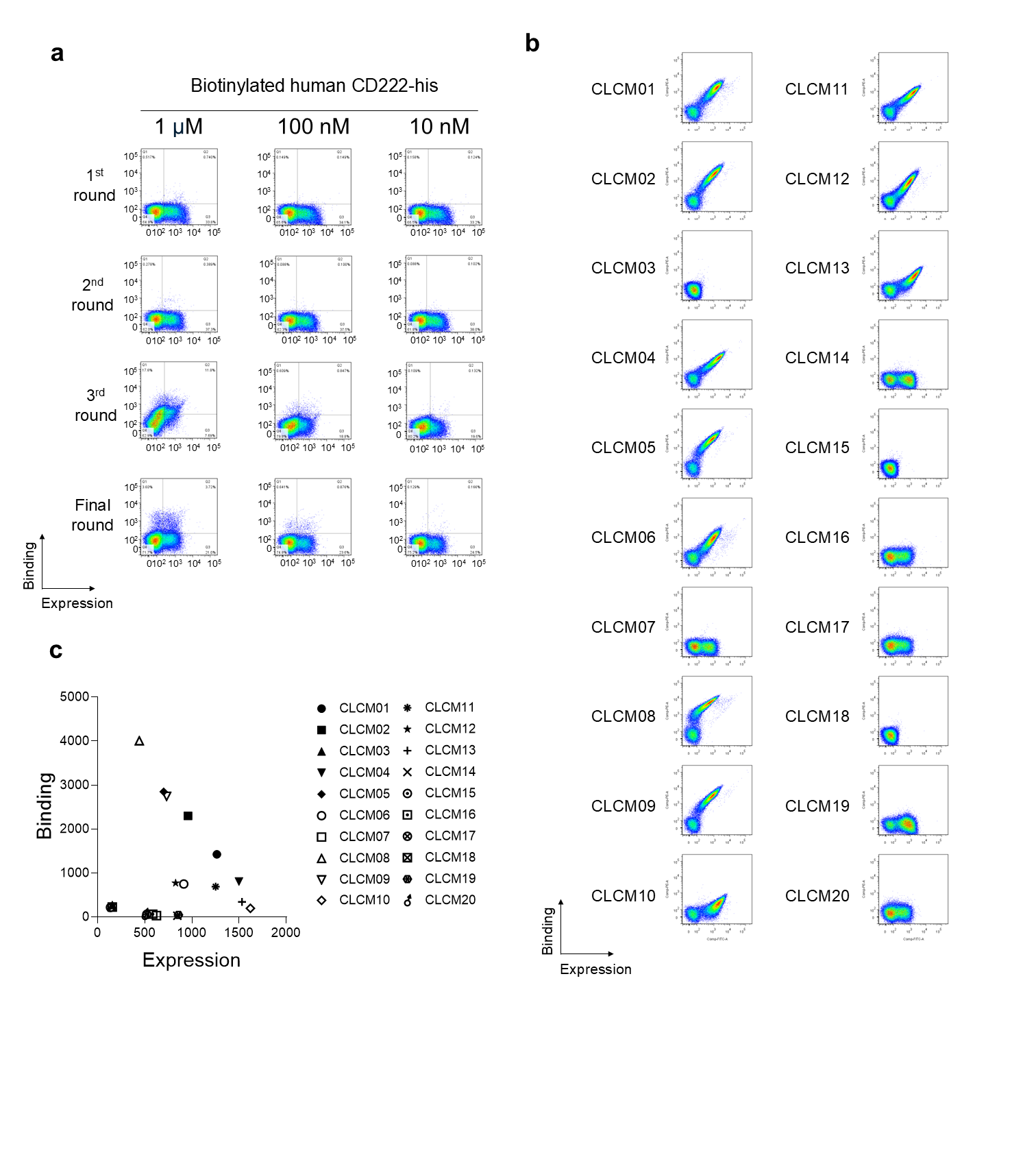

**Supplementary Figure 4. Sequential enrichment and clone-level analysis of CD222-binding yeast-displayed Fabs.**

(a) Representative flow-cytometric analysis of Fab expression and binding to biotinylated human CD222-His during sequential library-enrichment rounds. Binding was evaluated using 1 μM, 100 nM, and 10 nM biotinylated CD222-His as indicated, with progressive enrichment of CD222-binding populations across successive sorting rounds. (b) Representative flow-cytometric analysis of Fab expression and CD222 binding for 20 unique clones (CLCM01–CLCM20) isolated from the final enriched pool. (c) Scatter plot summarizing Fab-display level and CD222-binding signal for the 20 CLCM clones. Each point represents an individual clone, with Fab expression plotted on the x-axis and CD222 binding on the y-axis.

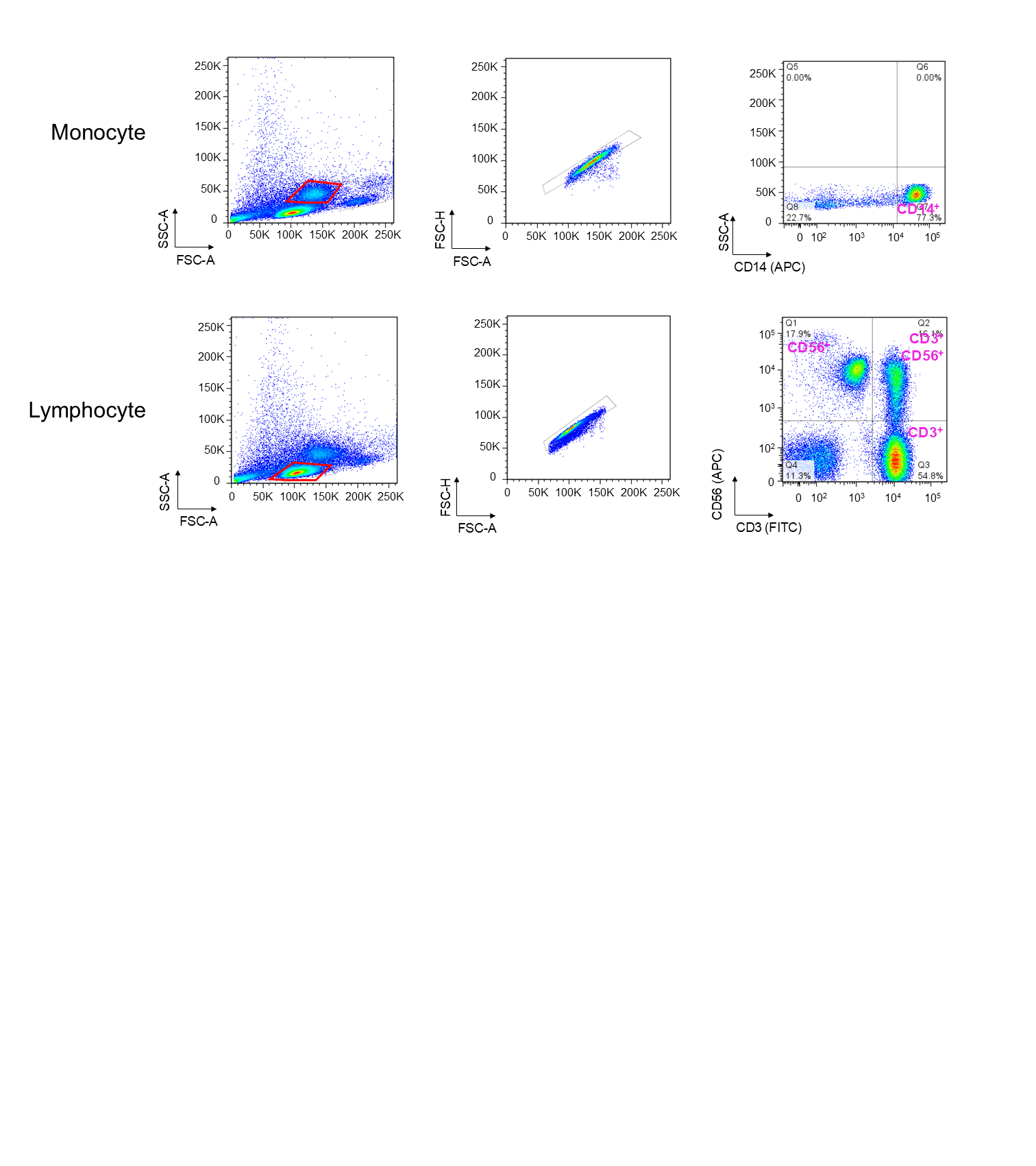

**Supplementary Figure 5. Flow-cytometric gating strategy for PBMC immune-cell subset analysis.**

Upper panels, monocytes were first identified from the FSC-A/SSC-A distribution, followed by singlet discrimination using FSC-H versus FSC-A, and subsequently gated as CD14-positive events. Lower panels, lymphocytes were first identified from the FSC-A/SSC-A distribution, followed by singlet discrimination using FSC-H versus FSC-A. CD3 and CD56 staining was then used to distinguish CD3-positive T cells, CD3-negative CD56-positive NK cells, and CD3-positive CD56-positive NKT-like cells. Monocyte and lymphocyte analyses were performed in separate staining panels.

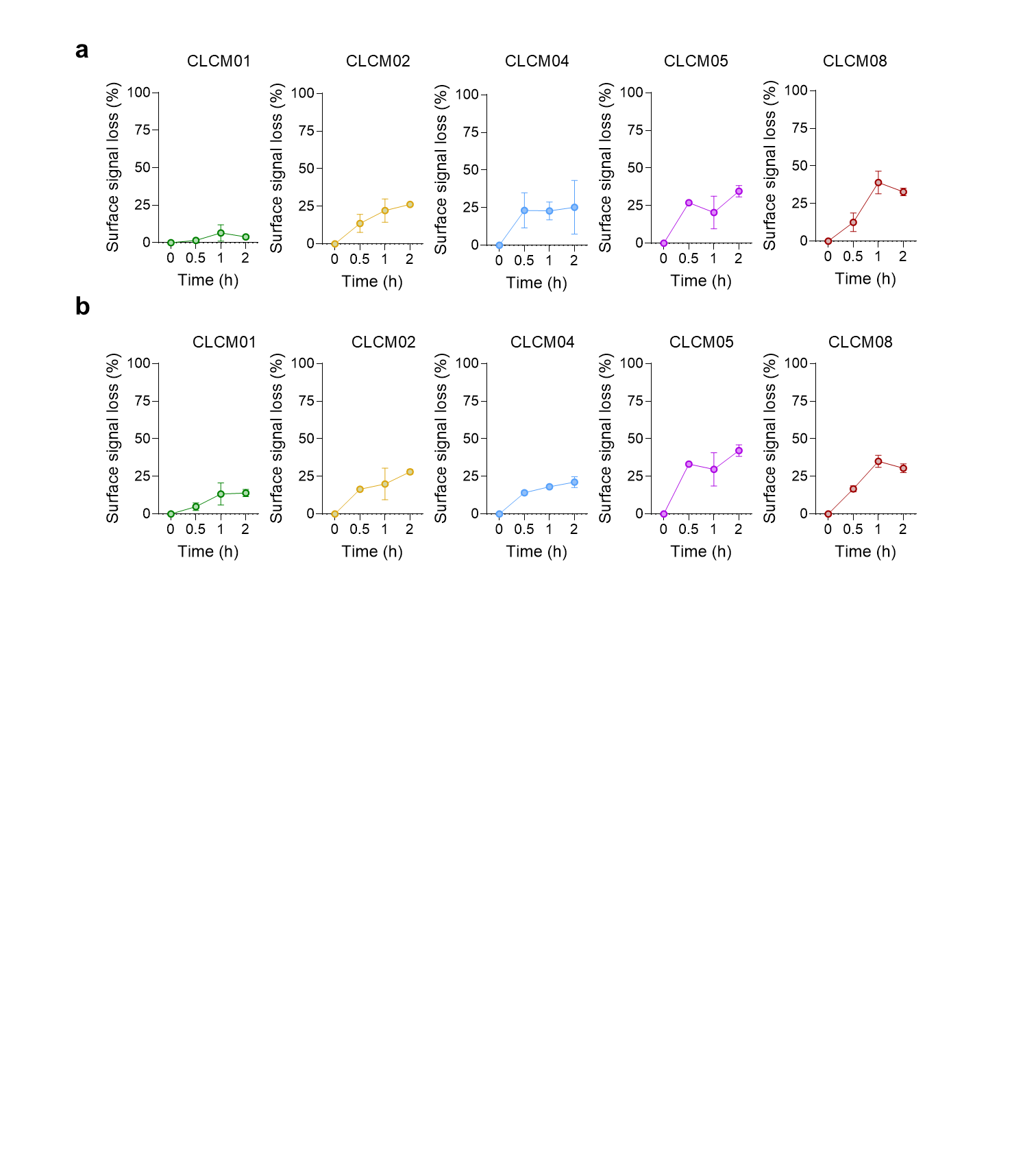

**Supplementary Figure 6. Temperature-dependent loss of surface-associated anti-CD222 antibody signal in primary immune cells.**

Primary PBMCs were labeled with the indicated biotinylated anti-CD222 antibodies at 4 °C, followed by detection with streptavidin-PE. Samples were subsequently maintained at 4 °C or incubated at 37 °C for 0, 0.5, 1, or 2 h. Temperature-dependent surface-signal loss was calculated at each time point as [1 − (MFI37 °C/MFI4 °C)] × 100 in (a) CD14-positive monocytes and (b) CD3-positive T cells. Because the assay did not distinguish internalized antibody from other potential causes of reduced surface-associated fluorescence, including antibody shedding or epitope masking, the readout was interpreted as temperature-dependent loss of surface-associated antibody signal rather than definitive antibody internalization. Data are presented as mean ± SD from two replicate measurements in a single donor, and no inferential statistical testing was applied.

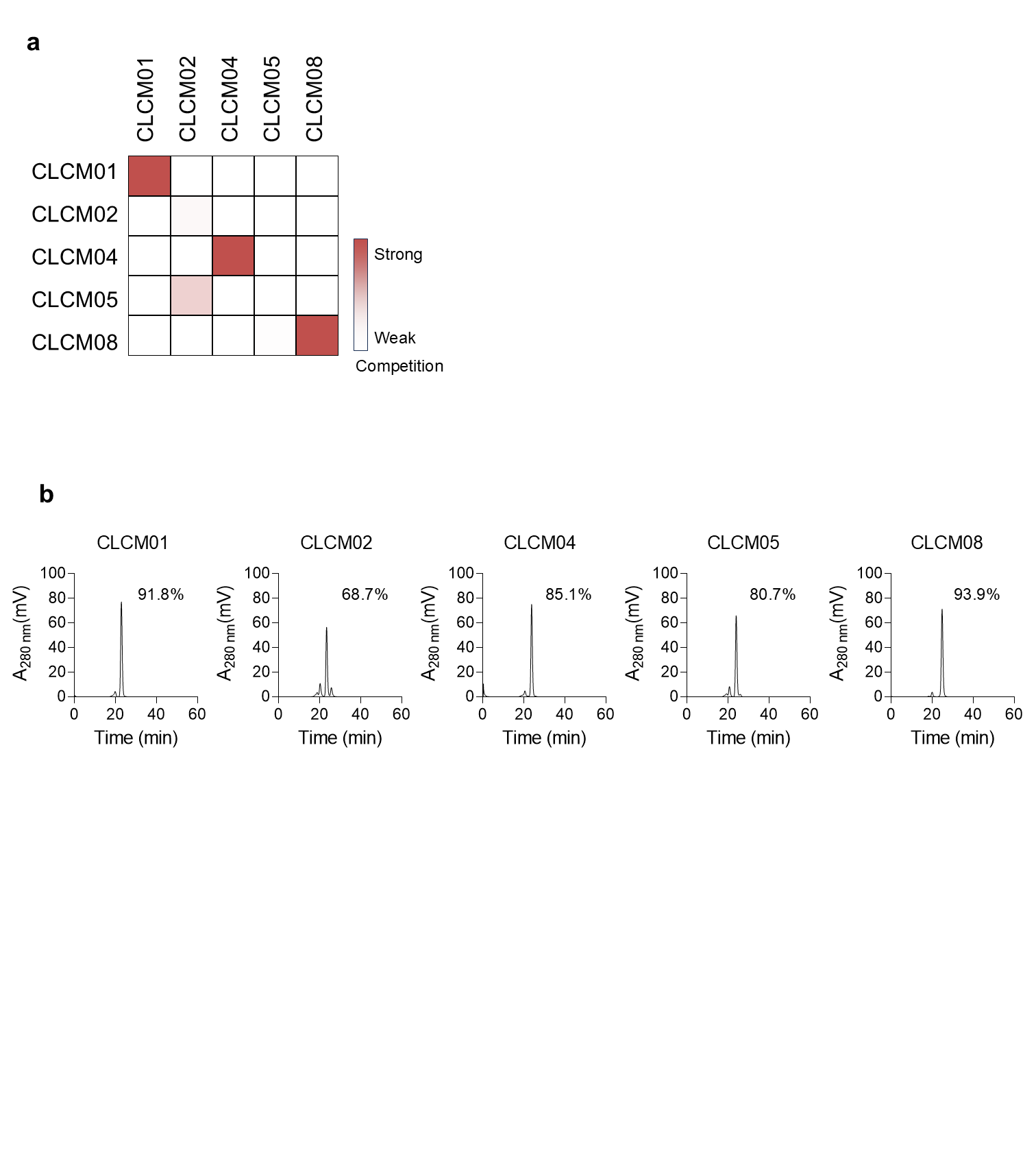

**Supplementary Figure 7. Competitive binding and SEC-HPLC characterization of selected anti-CD222 IgG clones.**

(a) Competitive ELISA analysis of the five selected anti-CD222 antibodies (CLCM01, CLCM02, CLCM04, CLCM05, and CLCM08). Immobilized anti-CD222 IgGs were incubated with biotinylated CD222-His in the presence of the indicated soluble competitor antibodies. Residual binding of biotinylated CD222-His was measured to assess pairwise competition among the antibody clones. The heatmap summarizes the relative degree of competition, with increasing color intensity indicating stronger competition. (b) SEC-HPLC chromatograms of the purified IgG clones. The percentages shown indicate monomeric content determined by peak-area integration: 91.8% for CLCM01, 68.7% for CLCM02, 85.1% for CLCM04, 80.7% for CLCM05, and 93.9% for CLCM08.

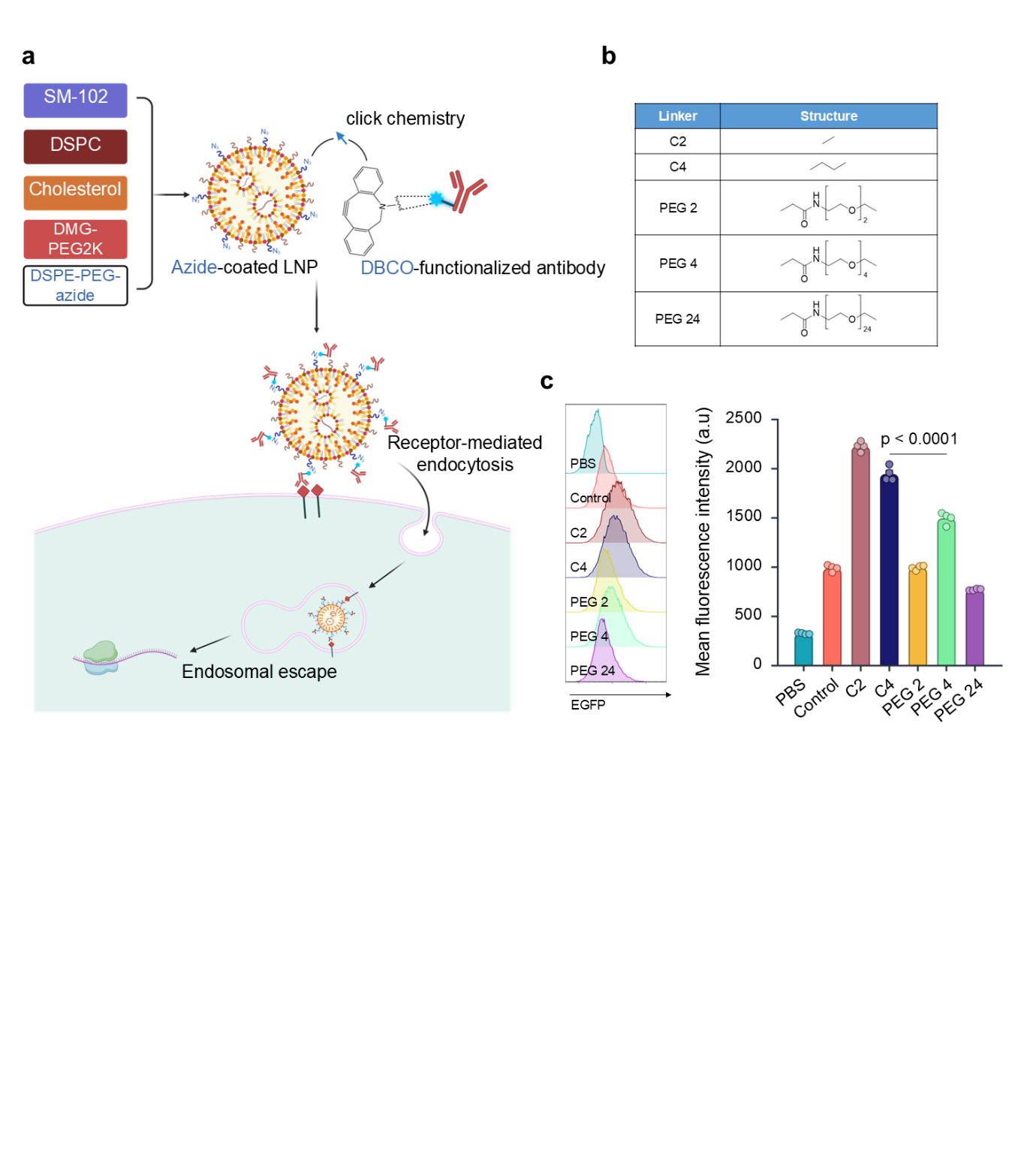

**Supplementary Figure 8. Optimization of antibody-LNP linker architecture for reporter mRNA delivery.**

(a) Schematic of antibody conjugation to azide-bearing SM-102 LNPs. DBCO-functionalized antibodies were coupled to azide-bearing LNPs by copper-free click chemistry, followed by cellular uptake and endosomal escape. (b) Chemical structures of the C2, C4, PEG2, PEG4, and PEG24 linkers evaluated for antibody conjugation. (c) Representative EGFP fluorescence distributions and corresponding mean fluorescence intensities in RAW 264.7 cells following treatment with LNP formulations prepared using the indicated linker chemistries. PBS and untargeted LNP were included as comparators. Among the tested linkers, the C2 linker produced the highest EGFP fluorescence intensity, reaching approximately 2,200 arbitrary units, and was therefore selected for subsequent antibody–LNP conjugate preparation. Data are presented as mean ± SD from four replicate measurements. Statistical significance was assessed by one-way ANOVA followed by Tukey’s multiple-comparison test. ****P < 0.0001.

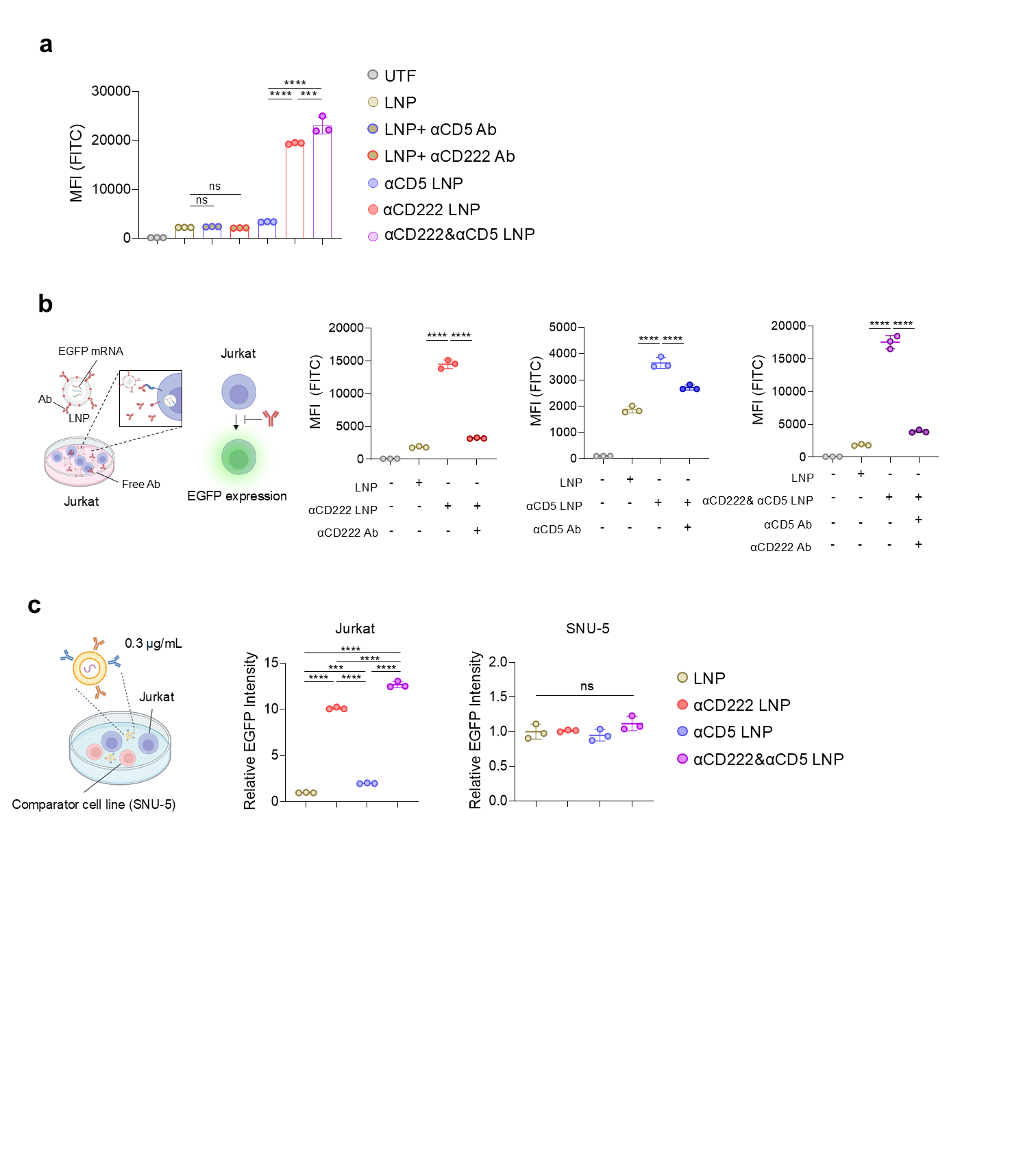

**Supplementary Figure 9. Validation of antibody-targeted EGFP mRNA-LNP delivery.**

(a) EGFP mRNA delivery by antibody-conjugated LNPs in Jurkat cells. Jurkat cells were treated with EGFP mRNA-loaded LNPs at 0.3 μg/mL, and EGFP expression was measured by flow cytometry. αCD222-LNP and dual αCD222/αCD5-LNP showed markedly enhanced EGFP delivery compared with untargeted LNP or free antibody-mixed controls. (b) Soluble-antibody competition analysis of targeted LNP delivery in Jurkat cells. αCD222-LNP and αCD5-LNP were tested in the absence or presence of excess soluble αCD222 or αCD5 antibody, respectively. Dual αCD222/αCD5-LNP was tested in the absence or presence of combined soluble αCD222 and αCD5 antibodies. Untargeted LNP was included as a reference condition. (c) Comparison of targeted EGFP mRNA delivery in Jurkat and SNU-5 cells at 0.3 μg/mL mRNA. EGFP fluorescence intensity was normalized to the corresponding untargeted-LNP condition within each cell line. Data are presented as mean ± SD from n = 3 independent experiments. Statistical significance was determined by one-way ANOVA followed by Tukey's multiple-comparison test. ns, not significant; *P < 0.05; **P < 0.01; ***P < 0.001; ****P < 0.0001. UTF, untransfected cells; LNP, untargeted LNP.

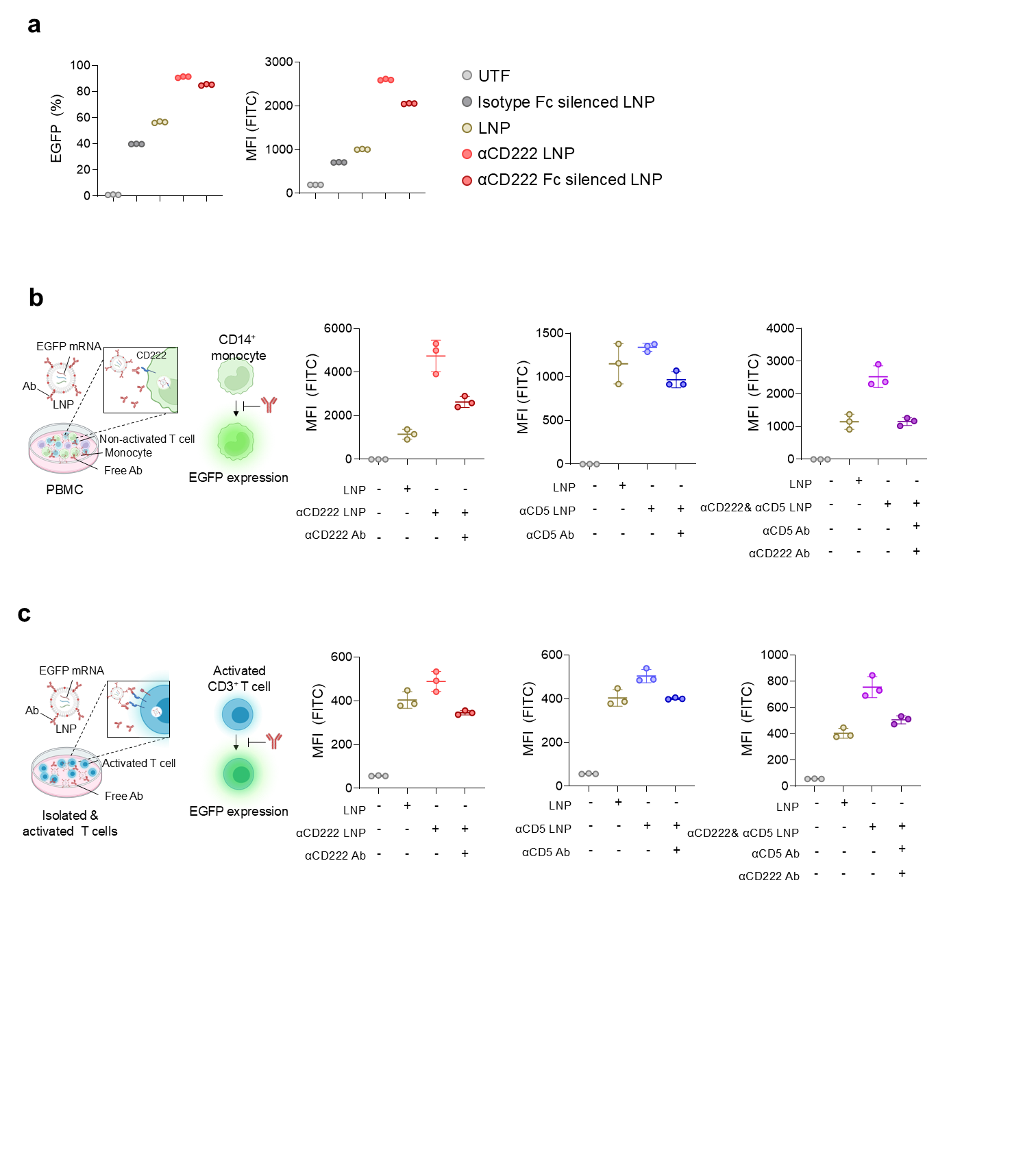

**Supplementary Figure 10. Assessment of Fc- and receptor-dependent contributions to targeted EGFP mRNA-LNP delivery**

(a) EGFP delivery in CD14-positive monocytes within PBMC cultures following treatment with cell-only control, Fc-silenced isotype-control LNP, untargeted LNP, αCD222-LNP, or Fc-silenced αCD222-LNP. EGFP-positive frequency and EGFP mean fluorescence intensity (MFI) were quantified by flow cytometry. (b) Soluble-antibody competition analysis in CD14-positive monocytes within PBMC cultures. PBMCs were treated with untargeted LNP, αCD222-LNP, αCD5-LNP, or dual αCD222/αCD5-LNP in the absence or presence of the corresponding excess soluble antibody or antibody combination, and EGFP MFI was quantified within the CD14-positive monocyte population. (c) Soluble-antibody competition analysis in isolated CD3/CD28-activated primary T cells using the same LNP and soluble-antibody combinations. EGFP MFI was quantified within the CD3-positive T-cell population.

Data are presented as mean ± SD of n = 3 technical replicates derived from a single donor. Comparisons are reported descriptively, and no inferential statistical testing was applied because technical replicates were not treated as independent biological observations. UTF, untransfected cells; LNP, untargeted LNP.

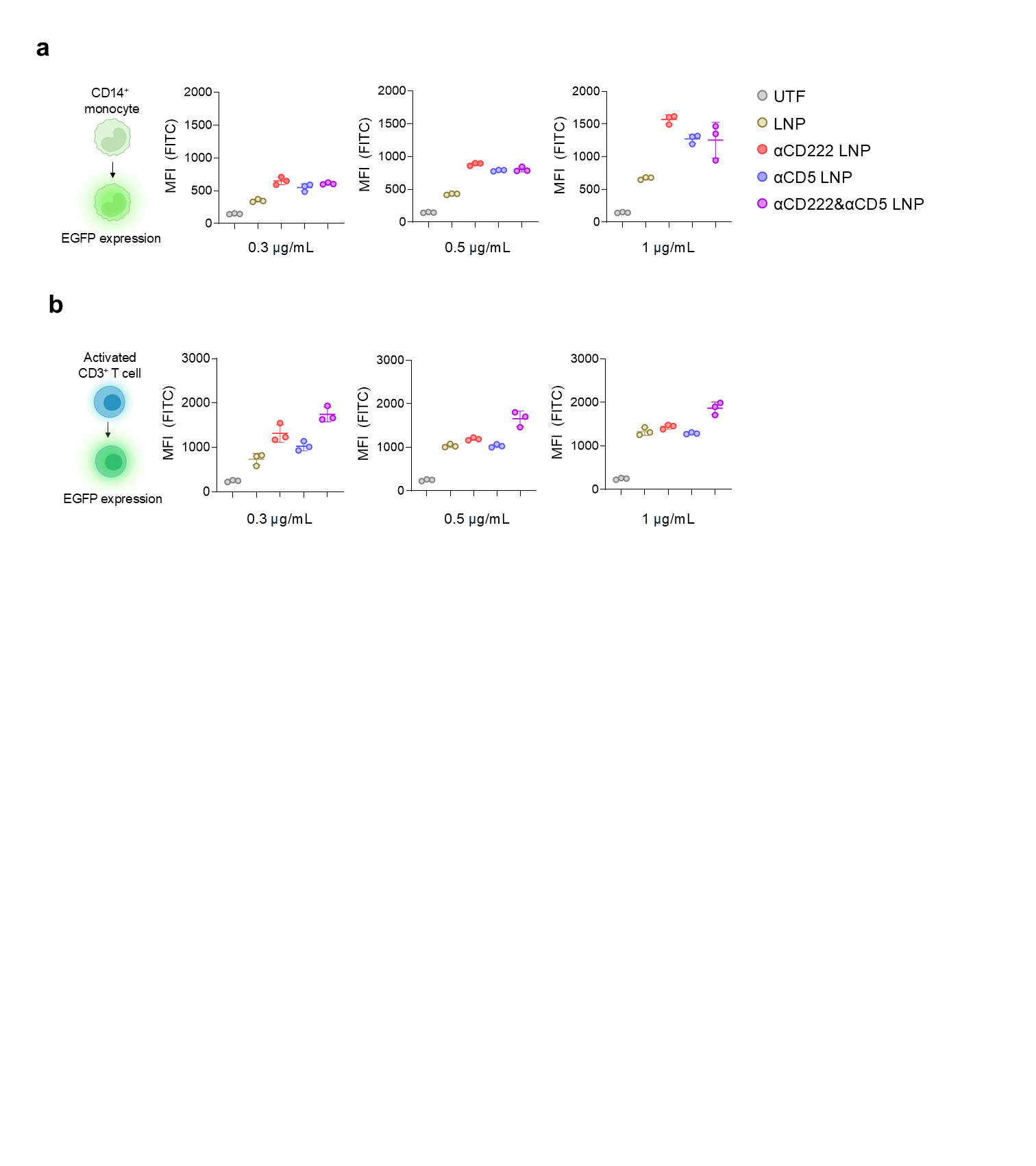

**Supplementary Figure 11. Dose-dependent EGFP expression intensity in primary monocytes and activated T cells.**

EGFP mean fluorescence intensity (MFI) was quantified by flow cytometry 24 h after treatment with the indicated EGFP mRNA-LNP formulations at final mRNA concentrations of 0.3, 0.5, or 1.0 μg/mL. (a) EGFP MFI in CD14-positive monocytes within non-activated PBMC cultures. (b) EGFP MFI in isolated CD3-positive T cells following 24 h of CD3/CD28 Dynabead activation before LNP treatment. Data are presented as mean ± SD of n = 3 technical replicates derived from a single donor. No inferential statistics were applied (Methods 2.24). UTF, untransfected cells; LNP, untargeted LNP.

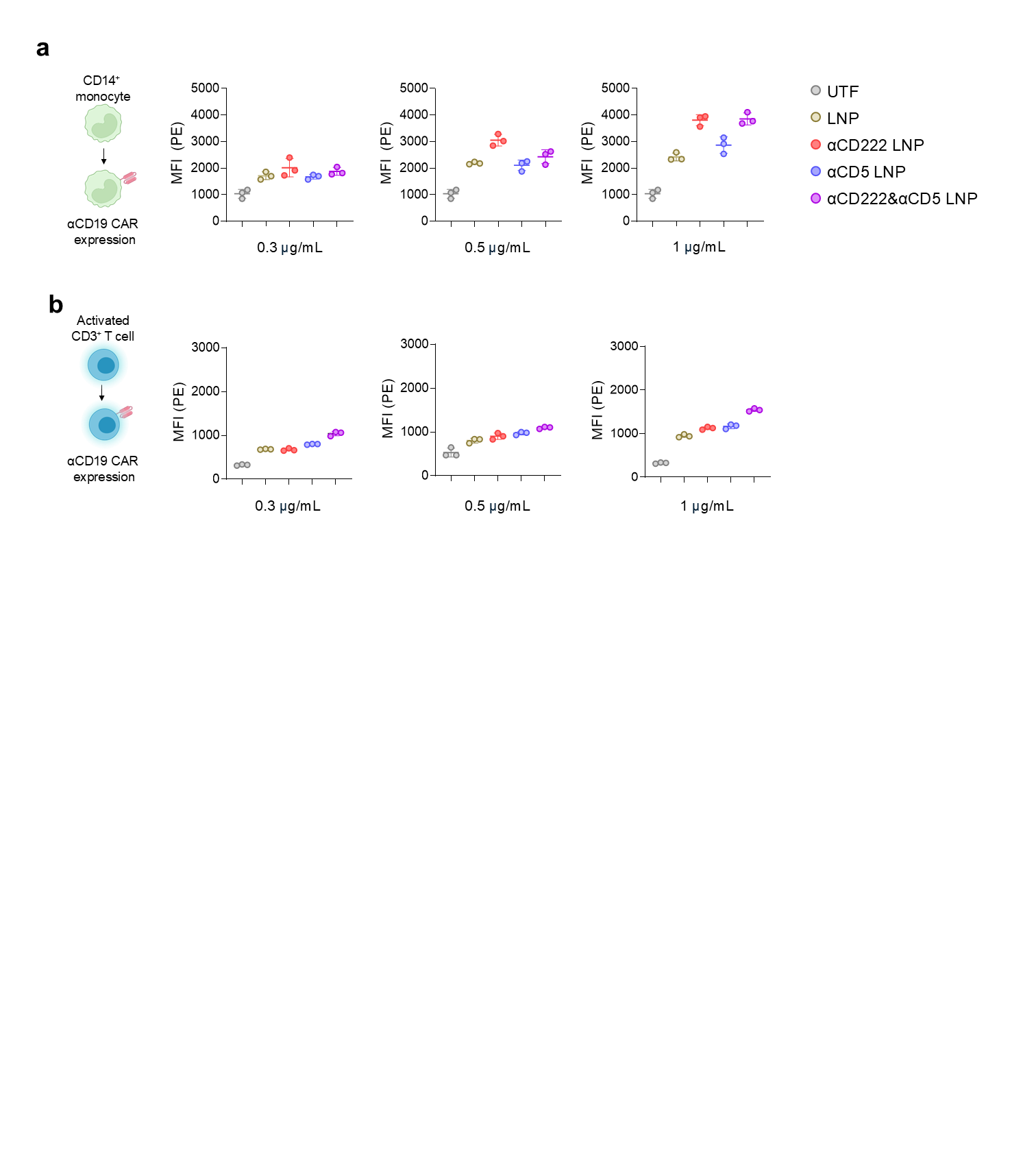

**Supplementary Figure 12. Dose-dependent CAR surface-expression intensity in primary monocytes and activated T cells.**

Surface expression of the anti-CD19 CAR was quantified by flow cytometry 24 h after treatment with the indicated CAR mRNA-LNP formulations at final mRNA concentrations of 0.3, 0.5, or 1.0 μg/mL. CAR expression was detected using a biotinylated anti-MYC antibody followed by streptavidin-PE, and PE mean fluorescence intensity (MFI) was quantified in (a) CD14-positive monocytes within non-activated PBMC cultures and (b) isolated CD3-positive T cells following 24 h of CD3/CD28 Dynabead activation before LNP treatment. Data are presented as mean ± SD of n = 3 technical replicates derived from a single donor. No inferential statistics were applied (Methods 2.24). UTF, untransfected cells; LNP, untargeted LNP.

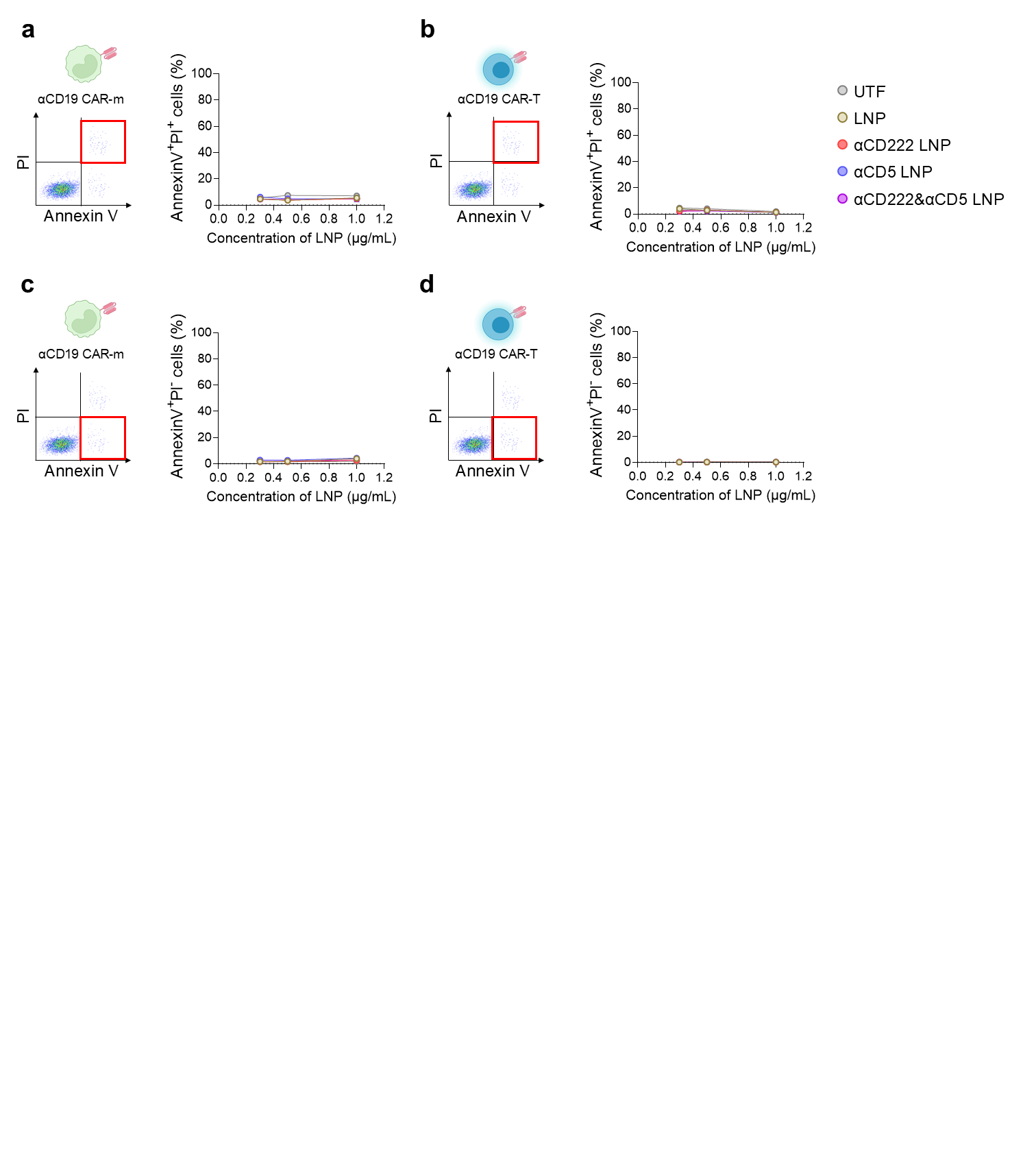

**Supplementary Figure 13. Apoptotic and membrane-compromised cell populations after CAR mRNA-LNP treatment.**

Annexin V/PI staining was performed 24 h after treatment with the indicated CAR mRNA-LNP formulations at final mRNA concentrations of 0.3, 0.5, or 1.0 μg/mL. Representative flow-cytometric gates and dose-dependent quantification are shown for (a) Annexin V-positive/PI-positive cells within the CD14-positive monocyte population of PBMC cultures, (b) Annexin V-positive/PI-positive cells within isolated CD3/CD28-activated T-cell cultures, (c) Annexin V-positive/PI-negative cells within the CD14-positive monocyte population of PBMC cultures, and (d) Annexin V-positive/PI-negative cells within isolated CD3/CD28-activated T-cell cultures. The corresponding Annexin V-negative/PI-negative viable-cell fractions are presented in the main figure. Data are presented as mean ± SD of n = 3 technical replicates derived from a single donor. No inferential statistics were applied (Methods 2.24). UTF, untransfected cells; LNP, untargeted LNP.

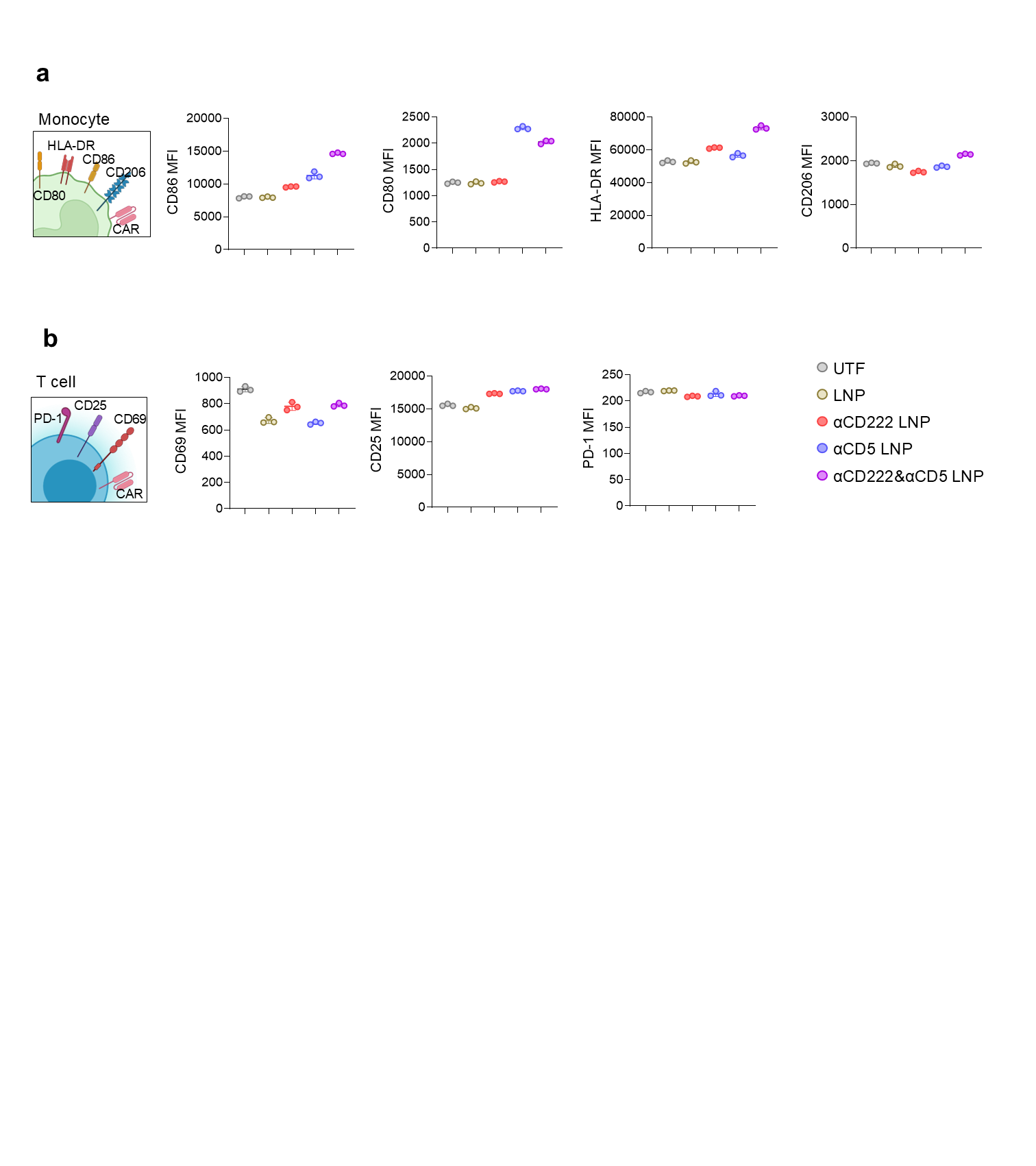

**Supplementary Figure 14. Absolute surface-marker expression after receptor-targeted CAR mRNA-LNP treatment.**

Absolute mean fluorescence intensity (MFI) values corresponding to the normalized surface-marker heatmaps presented in the main figure are shown. (a) Expression of CD86, CD80, HLA-DR, and CD206 in CD14-positive monocytes within non-activated PBMC cultures 24 h after treatment with the indicated CAR mRNA-LNP formulations. (b) Expression of CD69, CD25, and PD-1 in isolated CD3-positive T cells following CD3/CD28 activation and subsequent CAR mRNA-LNP treatment for 24 h. Data are presented as mean ± SD of n = 3 technical replicates derived from a single donor. No inferential statistics were applied (Methods 2.24). UTF, untransfected cells; LNP, untargeted LNP.

**Supplementary Table S1. Public single-cell RNA-sequencing datasets used to build the pan-cancer PBMC atlas.**

Dataset accession, repository, cancer or disease of origin, specimen and selected compartment, number of sample-level profiles, number of unique donors, library chemistry, sequencing platform, primary reference and source URL for each dataset included in the integrated atlas.

**Supplementary Table S2. Sample-level metadata for the integrated pan-cancer PBMC atlas.**

Dataset accession, sample accession, original sample name, donor or patient key, cancer or disease of origin, original specimen, library chemistry and source URL for every sample-level profile included in the atlas.

**Supplementary Table S3. Physicochemical characterization of the mRNA-LNP formulations used in this study.**

Hydrodynamic diameter, polydispersity index, zeta potential and mRNA encapsulation efficiency for untargeted, anti-CD222, anti-CD5 and dual anti-CD222/anti-CD5 mRNA-LNPs, together with the nominal antibody-input ratio of each formulation.
